# Structures of the PI3Kα/KRas complex on lipid bilayers reveal the molecular mechanism of PI3Kα activation

**DOI:** 10.1101/2025.03.22.644753

**Authors:** Hayarpi Torosyan, Michael D. Paul, Allison Maker, Brigitte G. Meyer, Natalia Jura, Kliment A. Verba

**Affiliations:** Cardiovascular Research Institute, University of California San Francisco; San Francisco, CA 94158, USA; Tetrad Graduate Program, University of California San Francisco; San Francisco, CA 94158, USA; Department of Cellular and Molecular Pharmacology, University of California San Francisco; San Francisco, CA 94158, USA; Quantitative Biosciences Institute (QBI), University of California San Francisco; San Francisco, CA 94158, USA

## Abstract

PI3Kα is a potent oncogene that converts PIP2 to PIP3 at the plasma membrane upon activation by receptor tyrosine kinases and Ras GTPases. In the absence of any structures of activated PI3Kα, the molecular details of its activation remain unknown. Here, we present cryo-EM structures of the PI3Kα/KRas complex embedded in lipid nanodiscs, revealing a rich ensemble of PI3Kα states adopted at the membrane surface. The sequential addition of a lipid bilayer, PIP2 and an activating phosphopeptide leads to the progressive release of key inhibitory domains from the PI3Kα catalytic core, which directly correlates with the reorganization of its active site. While association with POPC/POPS nanodiscs partially relieves PI3Kα autoinhibition, incorporation of PIP2 triggers near-complete displacement of PI3Kα inhibitory domains and significant restructuring of active site regulatory motifs. The addition of the activating phosphopeptide induces dimerization of the PI3Kα/KRas complex through a p110α catalytic subunit-mediated interface that is sterically occluded in autoinhibited PI3Kα. In cells, this dimeric PI3Kα complex amplifies Akt signaling in response to growth factor stimulation. Collectively, our structures map the conformational landscape of PI3Kα activation and reveal previously unexplored interfaces for potential therapeutic targeting.

## Results

Phosphoinositide 3-kinase α (PI3Kα) plays critical roles in normal growth and development by coordinating a broad range of essential cellular processes, including glucose absorption, anabolic metabolism, cell growth, and survival (*1-3*). Given its expansive role in signaling, PI3Kα activity is tightly regulated to ensure cellular homeostasis. In its basal state, PI3Kα exists as an autoinhibited heterodimer composed of a p85α regulatory subunit and a p110α catalytic subunit, in which p85α inhibits the catalytic activity of p110α **(Fig. 1A)** (*4*). During activation, PI3Kα integrates two main inputs: phosphorylated tyrosines, most commonly presented by activated receptor tyrosine kinases (RTKs), and GTP-bound Ras (*1*). RTKs and Ras engage p85α and p110α, respectively, recruiting PI3Kα to the membrane and relieving its autoinhibition. Once active, PI3Kα phosphorylates its substrate phosphatidylinositol 4,5-biphosphate (PIP2), generating phosphatidylinositol 3,4,5-biphosphate (PIP3), which activates downstream signaling, including the Akt pathway.

**Fig. 1.**
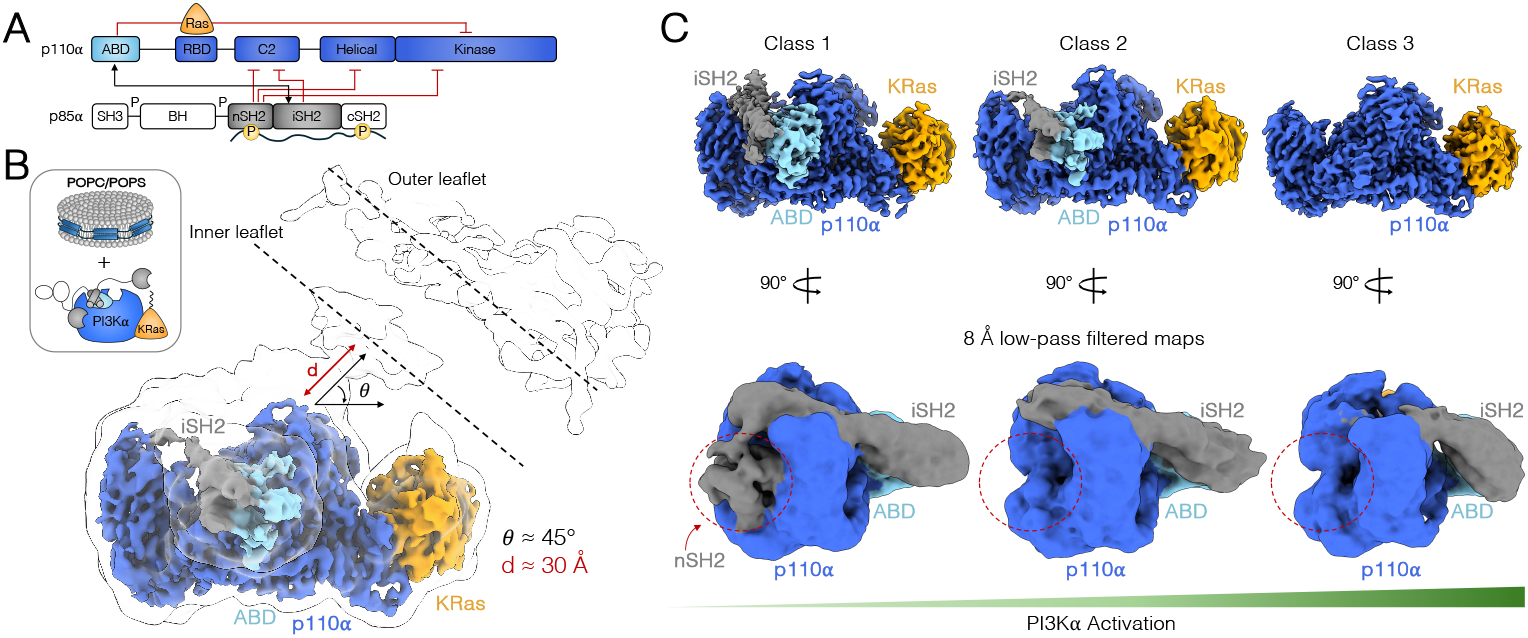
Structures of the PI3Kα/KRas complex bound to POPC/POPS nanodiscs reveal an increasingly activated PI3Kα. (**A**) Cartoon schematic of domain structure of p85α regulatory and p110α catalytic subunits of PI3Kα. p110α is composed of an adaptor binding domain (ABD), Ras binding domain (RBD), C2, helical, and kinase domains. p85α contains a Src homology 3 domain (SH3), BCR homology domain (BH), and two SH2 domains (nSH2 and cSH2) separated by a coiled coil domain, termed the inter-SH2 (iSH2) domain. Inter and intra-subunit inhibitory interactions are depicted with red lines and non-inhibitory interactions are illustrated with black arrows. (**B**) Cryo-EM map of the Class 2 PI3Kα/KRas complex bound to POPC/POPS nanodiscs overlayed with a transparent 10 Å low-pass filtered version of the same map to highlight membrane interface. p85α is colored in gray, p110α in blue, the ABD domain of p110α in cyan, and KRas in orange. (**C**) Cryo-EM maps of the three distinct classes of the PI3Kα/KRas complex bound to POPC/POPS nanodiscs as well as their 8 Å lowpass filtered counterparts, illustrating the dissociation of p85α from the p110α catalytic core.

Although activating oncogenic mutations occur in both PI3Kα subunits, the gene encoding p110α (*PIK3CA*), has emerged as one of the most frequently mutated genes in cancer (*5*). Oncogenic mutations significantly lower the activation threshold of PI3Kα by weakening inter and intra-subunit inhibitory contacts, often mimicking aspects of physiological PI3Kα activation. Even in the absence of activating mutations, PI3Kα plays a crucial role in promoting tumorigenesis driven by genetic lesions in upstream oncogenes such as Ras and RTKs, with the HER2/HER3 complex of the Human Epidermal Growth Factor Receptor (HER) family of RTKs being a primary driver (*6-9*). While the importance of synergistic interactions among RTKs, Ras, and PI3Kα is well-established in physiology and disease, their precise mechanistic basis remains unclear.

Extensive biophysical studies have broadly outlined global conformational changes that occur within PI3Kα in response to RTK-derived phosphopeptides and Ras (*10-15*). However, high-resolution understanding of PI3Kα activation—particularly in the context of a lipid bilayer where PI3Kα encounters its substrate, PIP2—has been limited by the dynamic nature of activated PI3Kα and its weak and transient association with synthetic membranes (*16-18*). Thus, while over 100 structures of PI3Kα have been solved to date, PI3Kα has yet to be structurally characterized in complex with either of its two activating inputs, RTKs and Ras, or its substrate, PIP2, on the surface of a lipid bilayer. To address these gaps in our understanding of PI3Kα biology, we used cryo-electron microscopy (cryo-EM) and lipid nanodisc-based reconstitution to investigate membrane-associated PI3Kα states in the presence of KRas, PIP2, and an activating phosphopeptide. Our work reveals the dynamic rearrangements undertaken by PI3Kα during activation at atomic resolution.

### POPC/POPS nanodiscs induce partial activation of PI3Kα

Reconstitution of PI3Kα alone on the surface of lipid nanodiscs did not yield stable complexes for high-resolution structural studies. Thus, to stabilize PI3Kα on the nanodisc surface, we incorporated GppNHp-loaded, farnesylated, and methylated KRas, modifications of KRas that mimic its processing in cells (*19*). We also included a molecular glue compound (D223), which binds at the PI3Kα/KRas interface, increasing PI3Kα’s affinity for KRas (*20*). Incubating purified PI3Kα with processed KRas, D223, and POPC/POPS lipid nanodiscs resulted in a stable complex suitable for cryo-EM (**fig. S1A-S1B)**. Analysis of this complex on graphene-oxide (GO) grids resulted in reconstructions of three distinct states at resolutions of 3.01 Å (class 1), 3.23 Å (class 2), and 3.05 Å (class 3) **(fig. S1C-S1K, fig. S2)**. In all three classes, the PI3Kα/KRas interaction is well resolved with the C-terminus of KRas reaching into the nanodisc **(Fig. 1B)**. The globular domain of KRas is positioned far from the nanodisc surface, lacking secondary interactions with membrane lipids beyond the farnesylated C-terminus. Although the resolution at the KRas- nanodisc interface (KRas residues 173-189) does not allow for *de novo* model building, we observe density for the residues connecting with the membrane inner leaflet in the low-pass filtered maps **(fig. S3A)**. PI3Kα interacts with KRas via its Ras Binding Domain (RBD) in a similar manner to that of the recently published p110α/KRas crystal structure, which is also stabilized by a molecular glue (*20*). Although the KRas-RBD domain conformations are essentially invariant between our structure and the recently published work (RMSD 0.661Å) **(fig. S3B)**, there is a visible rotation in the kinase domain between the two structures, either due to the presence of p85α or the lipid bilayer (**fig. S3C**). In both structures, KRas forms extensive interactions with the p110α RBD via its switch I and switch II regions (**fig. S3D**), but makes no contacts with the p110α kinase domain, in contrast to the previously solved p110γ/HRas structure (*21*) **(fig. S3E**).

Global examination of the three POPC/POPS-bound classes of the PI3Kα/KRas complex reveal increasingly activated states of the PI3Kα holoenzyme. This is reflected in the gradual destabilization of the inhibitory interfaces formed by the nSH2 and inter-SH2 (iSH2) domains of p85α, which wrap around p110α in the absence of activating inputs, inhibiting its kinase activity (*4,22,23*). Part of this inhibition involves the stabilization of an intra-p110α subunit interface between the kinase domain (KD) and the Adapter Binding Domain (ABD), which gradually unravels across the three classes **(Fig. 1C)**.

In class 1, the inhibitory iSH2 and ABD domains are well resolved along with the remainder of p110α. The nSH2 domain is present, although at a partial occupancy, visible only in an 8 Å low-pass filtered map **(Fig. 1C, left)**. Nevertheless, this most inhibited state is still in a more active conformation than observed in any of the previously published structures of wild-type (WT) PI3Kα in which the nSH2 and iSH2 domains are well-ordered **(table S1)**. Class 2 adopts an intermediate conformation with no observable p110α-nSH2 contacts and increasingly dynamic interactions between p110α and iSH2 and ABD domains **(Fig. 1C, middle)**. Finally, in class 3, the most active-like state in this dataset, all three inhibitory interfaces involving the nSH2, iSH2, and ABD domains are disrupted, with the iSH2 and ABD domains only partially resolved in low-pass filtered maps **(Fig. 1C, right)**.

The release of inhibitory interactions is coupled with the rearrangement of key regulatory motifs within the active site of PI3Kα, communicated through the separation of the C2 and kinase domains **(fig. S4A)**. The kinase activation loop, which in the class 1 reconstruction is stabilized in the collapsed, inactive conformation primarily by the iSH2 domain, becomes more dynamic and poorly resolved in class 2 and class 3 states **(fig. S4B)**. Additionally, residues corresponding to helix kα12 of the p110α regulatory arch fold as a coil away from the active site and are situated between helices kα8 and kα11 in the class 1 state, as seen in published crystal structures of autoinhibited PI3Kα (*4*,*24*,*25*). However, as PI3Kα adopts more active-like conformations in classes 2 and 3, helix kα12 residues become increasingly dynamic and poorly resolved **(fig. S4C)**. These conformational changes are likely a result of PI3Kα-membrane interactions, and not Ras binding. In our class 1 structure, PI3Kα adopts a more active-like conformation compared to the crystal structure of the p110α/KRas complex, in which the entire p85α subunit is absent **(fig. S3C)**. This suggests that the primary role of Ras is to anchor PI3Kα to the membrane, rather than directly induce an activating conformational change. Instead, membrane binding appears to be critical to triggering both global and local conformational changes in PI3Kα, consistent with its partial activation **(fig. S4D)**.

Consequently, the conformational rearrangements observed across the three classes translate to changes in PI3Kα-membrane interactions. In class 1, the p110α active site is ∼30Å away from the nanodisc and at a 45° angle relative to the lipid plane, while classes 2 and 3 show progressively closer interactions between PI3Kα and the nanodisc **(fig. S5A)**. However, even in class 3, the active site remains far from the lipid interface, precluding binding of a lipid substrate. This unproductive kinase/substrate conformation may result from the additional density that separates the p110α active site and the nanodisc inner leaflet, which is only resolved at low resolution. Remarkably, docking an AlphaFold3 model of full-length PI3Kα and flexibly fitting it into a 10 Å low-pass filtered class 1 map places the cSH2 domain of p85α within this region. **(fig. S5B)**. The elusive cSH2 domain of PI3Kα has never been structurally characterized within the context of the holoenzyme. Its membrane-proximal location in our structure contrasts with its position in the previously solved solution crystal structure of PI3Kβ, but is consistent with biophysical studies showing its propensity to bind lipids (*26-29*) **(fig. S5C)**.

### PIP2-containing nanodiscs engage and reorganize the active site of PI3Kα

Within the heterogeneous lipid bilayer, PI3Kα specifically phosphorylates PIP2 (*30*). However, previous structural studies of PI3Kα/PIP2 binding have been limited to the use of soluble PIP2 mimetics, like diC4-PIP2 (*31*), which do not fully recapitulate the behavior of the native substrate. We reconstituted the PI3Kα/KRas complex, as described above, on POPC/POPS nanodiscs containing 5% PIP2 **(fig. S6A, S6B)** and determined its structure by cryo-EM on PEG-amino GO grids at 2.82 Å resolution **(fig. S6C-S6G, fig. S7)**. Remarkably, the addition of PIP2 significantly reoriented PI3Kα relative to the membrane, increasing the buried surface area at the membrane interface and positioning the active site of PI3Kα in direct contact with membrane lipids **(Fig. 2A)**.

**Fig. 2.**
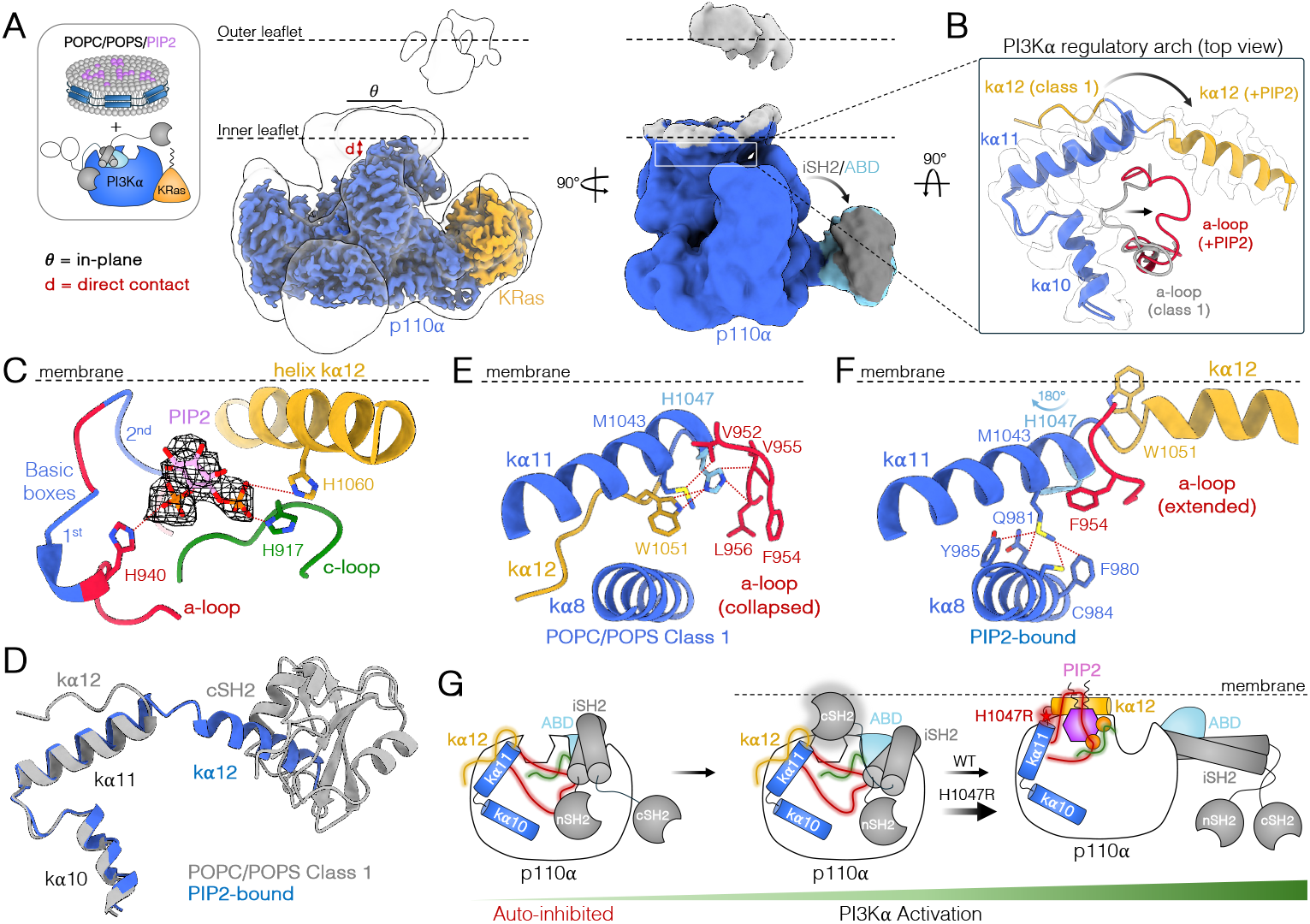
PIP2-containing nanodiscs reorganize the active site of PI3Kα. (**A**) Cryo-EM map of the PI3Kα/KRas complex bound to POPC/POPS/PIP2 nanodiscs overlayed with a transparent 10 Å low-pass filtered version of the same map (left) and an 8 Å low-pass filtered map (right). p110α is shown in blue, ABD of p110α in cyan, iSH2 in gray, and KRas in orange. (**B**) POPC/POPS-bound Class 1 and POPC/POPS/PIP2-bound regulatory arches overlayed with a transparent 4 Å low-pass filtered cryo-EM density of the PIP2-bound regulatory arch. kα10 and kα11 helices are shown in blue, kα12 in yellow, POPC/POPS Class 1 activation loop (a-loop) in gray, and PIP2-bound a-loop in red. (**C**) PIP2 headgroup (pink) overlayed with 4 Å low-pass filtered density (black) within the active site of PI3Kα, demonstrating its interaction with the a- loop, catalytic loop (c-loop, green), and helix kα12. The fully modeled activation loop is from the 5 Å low-pass filtered model of the PIP2-bound complex. Activation loop basic boxes (1^st^: residues 941-944, “KKKK”; 2^nd^: residues 948-951, “KRER”) are colored in blue. (**D**) Overlayed PI3Kα regulatory arches of the POPC/POPS-bound class 1 (gray) and PIP2-bound (blue) states, illustrating the steric clash between the cSH2 domain in the POPC/POPS-bound class 1 structure with the active conformation of helix kα12 in the PIP2-bound structure. (**E-F**) POPC/POPS class 1 and PIP2-bound active sites illustrating (E) the stabilization of helix kα12 and the a-loop in their inactive conformations by helices kα8 and kα11 in the POPC/POPS class 1 state and (F) the release of these inhibitory interactions in the PIP2-bound state. H1047 is highlighted in light blue. (**G**) Cartoon schematic illustrating the rearrangement of active site regulatory motifs upon activation of PI3Kα by POPC/POPS and PIP2-containing membranes. The location of the H1047R mutation is depicted with a red star.

In addition to expanding the PI3Kα/membrane interface, PIP2 induced both global and local conformational rearrangements in PI3Kα. Globally, the inhibitory iSH2 and ABD domains, which are partially displaced upon binding POPC/POPS membranes, completely disengage from the catalytic core of p110α in the presence of PIP2 **(Fig. 2A)**. Locally, the active site of PI3Kα also reorganizes to accommodate PIP2 binding **(fig. S9A)**. Specifically, the activation loop extends towards the membrane, while helix kα12 residues undergo a ∼180° rotation towards the active site, relative to the POPC/POPS class 1 state, and adopt a defined helical structure **(Fig. 2B)**. The inwardly rotated helix kα12 is parallel to the membrane plane and directly interacts with membrane lipids, serving as an additional anchor. Exhaustive structural analysis of this key regulatory motif across all structures of Class I PI3K enzymes confirmed that this conformation of helix kα12 is novel and likely induced by the presence of PIP2 (**fig. S8A**).

In their newly adopted conformations, helix kα12 and the activation loop wedge PIP2 between them, trapping it within the active site for phosphorylation **(Fig. 2C)**. We observe density for the PIP2 headgroup, with its 4’ and 5’ phosphates coordinated by helix kα12 and the activation loop, respectively. This is consistent with previous biochemical data, which showed that helix kα12 plays a critical role in PI3Kα membrane binding and catalytic activity (*25*). A conserved feature of helix kα12, called the “WIF” motif (W1057, I1058, F1059), is central to these functions. In our structure, W1057 engages the catalytic loop and I1058 contributes to interactions with a neighboring loop (p110α residues 868-874) at the membrane/protein interface **(fig. S9B, S9C)**. In its extended state, the highly basic activation loop reaches towards the membrane; however, the resolution for this region (residues 941-950) is too low to visualize specific sidechain interactions. By fitting only the backbone of this region into a 5 Å low-pass filtered map, we showcase the proximity of its 1^st^ and 2^nd^ basic boxes to the anionic membrane interface **(Fig. 2C)**. Notably, the position of the cSH2 domain in the structure of the complex obtained without PIP2 (POPC/POPS- class 1) directly clashes with the inward conformation of helix kα12 **(Fig. 2D)**. This suggests a potential role for the cSH2 domain in blocking non-specific membrane association of helix kα12 and subsequent PI3Kα activation.

### H1047R activates PI3Kα by destabilizing the inactive conformation of helix kα12

Our POPC/POPS and PIP2-bound structures offer mechanistic insights into the most common PI3Kα oncogenic mutation, H1047R, which has remained poorly understood. In the most inactive conformation, observed in the POPC/POPS class 1 state, H1047 faces the active site and forms hydrophobic contacts with both helix kα12 residues (folded back as a coil) and the C-terminus of the activation loop, stabilizing their inactive conformations **(Fig. 2E, fig. S9D)**. Upon PIP2 binding, H1047 flips ∼180°, resulting in the release of both inhibitory interactions, allowing the activation loop and helix kα12 to adopt their respective active conformations **(Fig. 2F)**. An introduction of a bulky, positively charged sidechain at this position, like the H1047R mutation, is predicted to disrupt the inhibitory hydrophobic interactions centered on H1047, destabilizing the inactive state of PI3Kα and facilitating the inward rotation and restructuring of helix kα12. This mechanism is supported by the observation that published structures of PI3Kα in which helix kα12 residues are found in an inward conformation are predominantly those of the H1047R mutant **(fig. S8B, S8C)** (*32*). However, in these structures, kα12 residues never adopt a helical structure, likely due to the absence of a membrane. Collectively, our structures suggest that upon recruitment of PI3Kα to the membrane, helix kα12’s affinity for membrane lipids drives its release from helices kα8 and kα11 and triggers a conformational switch which allows it to anchor PI3Kα to the membrane and stabilize the PIP2-bound state. H1047R reduces the energy barrier for this conformational change in helix kα12, resulting in the hyperactivation of PI3Kα **(Fig. 2G)**.

### Phosphopeptide binding induces dimerization of the PI3Kα/KRas complex

An essential step in PI3Kα membrane recruitment and activation is the binding of p85α’s tandem SH2 domains to phosphorylated tyrosines within YxxM motifs of PI3Kα activators, such as RTKs (*23*). To investigate how YxxM motifs cooperate with Ras to activate PI3Kα at the membrane, we reconstituted the PI3Kα/KRas complex on PIP2-containing lipid nanodiscs in the presence of a YxxM bis-phosphopeptide derived from HER3 **(fig. S10A, S10B)**. In the 2.89 Å cryo-EM reconstruction of the peptide-bound PI3Kα/KRas complex, PI3Kα is in an active conformation and engaged in extensive direct membrane interactions, similar to the PIP2-bound reconstruction in the absence of peptide **(Fig. 3A; fig. S10C-S10I, fig. S11)**. The density for the p85α subunit is not observed, and the ABD domain undergoes additional rotation away from the catalytic core of p110α relative to the PIP2-only bound state (**fig. S12A)**.

**Fig. 3.**
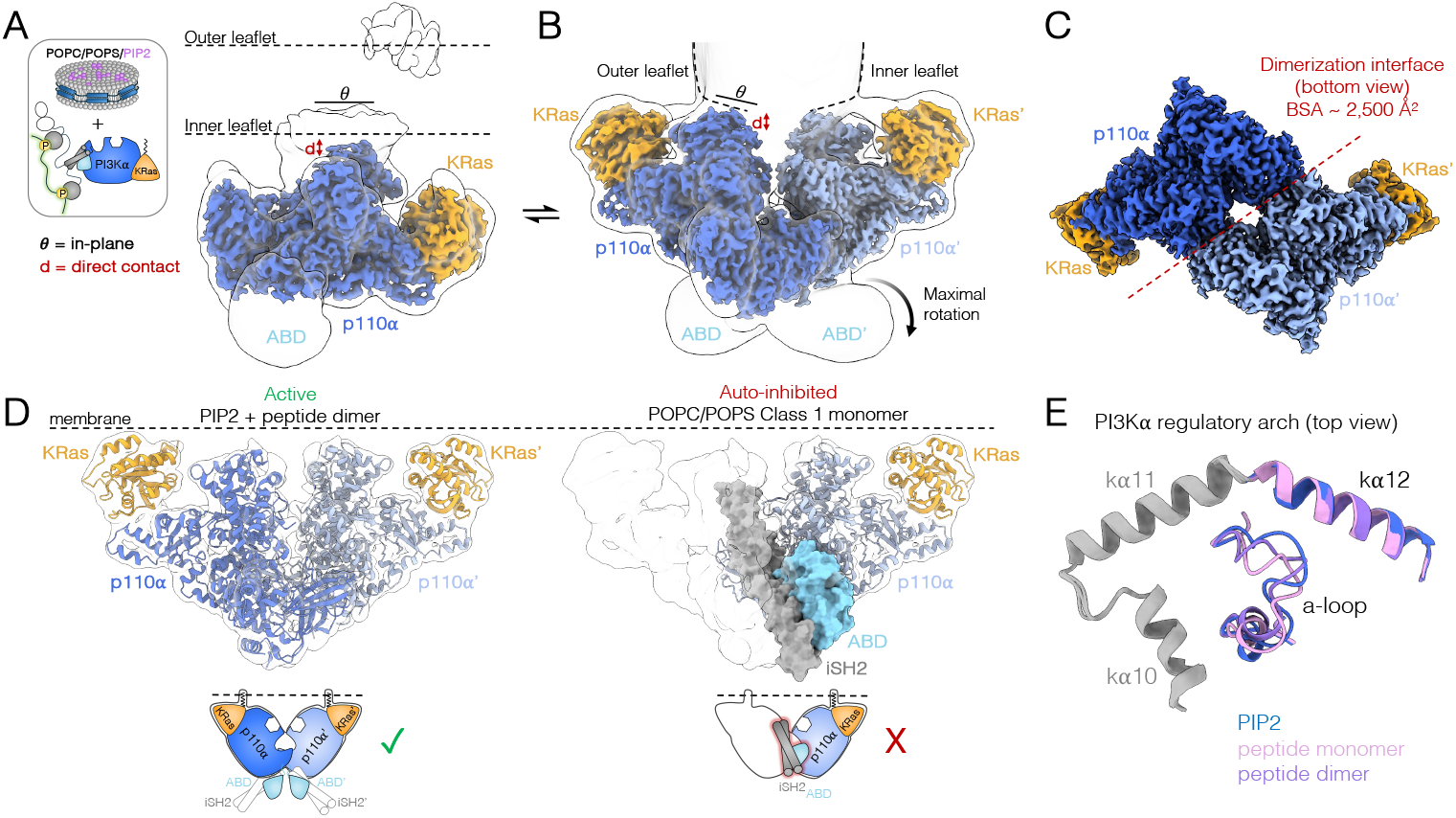
Phosphopeptide binding induces dimerization of the PI3Kα/KRas complex. (**A**) Cryo-EM map of the active monomeric PI3Kα/KRas/phosphopeptide complex bound to POPC/POPS/PIP2 nanodiscs overlayed with a transparent 10 Å low-pass filtered version of the same map. p110α is shown in blue and KRas in orange. (**B**) Cryo-EM map of the active dimeric PI3Kα/KRas/phosphopeptide complex bound to POPC/POPS/PIP2 nanodiscs overlayed with a transparent 10 Å low-pass filtered version of the same map. The p110α subunits are colored in light and dark blue and KRas molecules are colored in orange. (**C**) Bottom-up view of the dimer cryo-EM map highlighting the extensive dimerization interface of the PI3Kα/KRas/phosphopeptide complex. (**D**) Structure of the dimeric complex overlayed with its 8 Å low-pass filtered cryo-EM map (left) and the POPC/POPS-bound class 1 complex fit into the same map (right), illustrating the steric clash between the inhibitory iSH2 and ABD domains in their autoinhibited conformations (shown in surface representation) and the dimerization interface. (**E**) Top-down view of overlayed PI3Kα regulatory arches of the PIP2-bound (blue) and the PIP2/phosphopeptide-bound (monomeric [pink] and dimeric [purple]) structures highlighting the active conformations of helix kα12 and the activation loop.

Strikingly, a subset of particles in this dataset corresponded to a significantly larger complex oriented at the surface of the lipid nanodisc compared to datasets obtained without the peptide. Analysis of these particles yielded a 2.61 Å reconstruction of a previously uncharacterized dimeric PI3Kα/KRas complex **(Fig. 3B; fig. S10C-S10I)**, in which the density for p85α is no longer present. The extensive dimerization interface (BSA = 2,568 Å^2^) is mediated by the two p110α subunits **(Fig. 3C)** and is accompanied by maximal displacement of the ABD domain away from the catalytic core of p110α, which would otherwise sterically block the formation of the dimer **(Fig. 3D, fig. S12B)**. Thus, we infer that the peptide-induced PI3Kα/KRas dimer represents the most active PI3Kα state observed to date.

In the PI3Kα/KRas dimeric structure, each PI3Kα/KRas pair makes membrane interactions with opposite sides of the nanodisc **(Fig. 3B)**. This peculiar architecture may arise from the limited membrane surface area provided by the lipid nanodisc and the need to accommodate a complex as large as the PI3Kα/KRas dimer. In this context, PI3Kα balances its affinity for membrane lipids with its propensity for dimerization, taking advantage of the flexible linkage of KRas to the nanodisc to anchor the complex on both sides. We analyzed all nanodisc-bound structures of Ras alone deposited in the Protein Data Bank, and observed considerable variability in Ras insertion angles across the 13 NMR ensembles, underscoring Ras’s ability to dynamically engage with the membrane and adapt to different membrane curvatures **(fig. S13A, S13B)**. We propose that this flexibility at the Ras/membrane linkage enables the PI3Kα/KRas dimer to bind to the same cell membrane leaflet, without inducing significant membrane curvature.

The regulatory motifs in the PI3Kα active site adopt nearly identical conformations in the monomeric and dimeric phosphopeptide-bound PI3Kα/KRas complexes as seen in the PIP2-only bound state. In all three structures, the activation loop is extended, while helix kα12 of the regulatory arch is helical in nature and adopts an inward conformation **(Fig. 3E)**. We do not observe PIP2 density in the phosphopeptide-bound reconstructions and the overall resolution of the active site is lower. For the monomer, this may be attributed to the lower number of particles in the resulting reconstruction (111,698 particles), compared to the PIP2-only reconstruction (223,376 particles). In the case of the dimer, this is likely a consequence of the limited membrane surface area available to the active site in combination with the likelihood of encountering PIP2 within that space.

### PI3Kα homodimers amplify Akt signaling in cells

To investigate the physiological relevance of the p110α-mediated PI3Kα dimer, we first tested if PI3Kα forms homodimers in cells. The PI3Kα homodimer interface is largely stabilized by polar contacts between the C2 domain of one p110α subunit and the ABD-RBD linker and kinase domain of the second p110α subunit **(Fig. 4A, 4B)**. A key salt-bridge between R349 of the C2 domain and E707 of the kinase domain is central to this interface (**Fig. 4B**). Flag-tagged and HA-tagged WT PI3Kα were first co-expressed in MCF7 cells, and their interaction was tested by co-immunoprecipitation under serum-starved and neuregulin (NRG1β)-stimulated conditions to induce HER3 phosphorylation. We observed robust co-immunoprecipitation of the two differently tagged PI3Kα variants upon NRG1β stimulation (**Fig. 4C, 4D)**. However, NRG1β-induced PI3Kα dimerization was significantly diminished by mutations designed to break the p110α-mediated homodimer interface by inducing a charge flip in the R349-E707 salt-bridge (R349E, E707R) (**Fig. 4B-4D**). A mutation targeting weaker packing interactions at the interface (P124A), had no effect, indicating the overall robustness of the interface. (**Fig. 4C, 4D**).

**Fig. 4.**
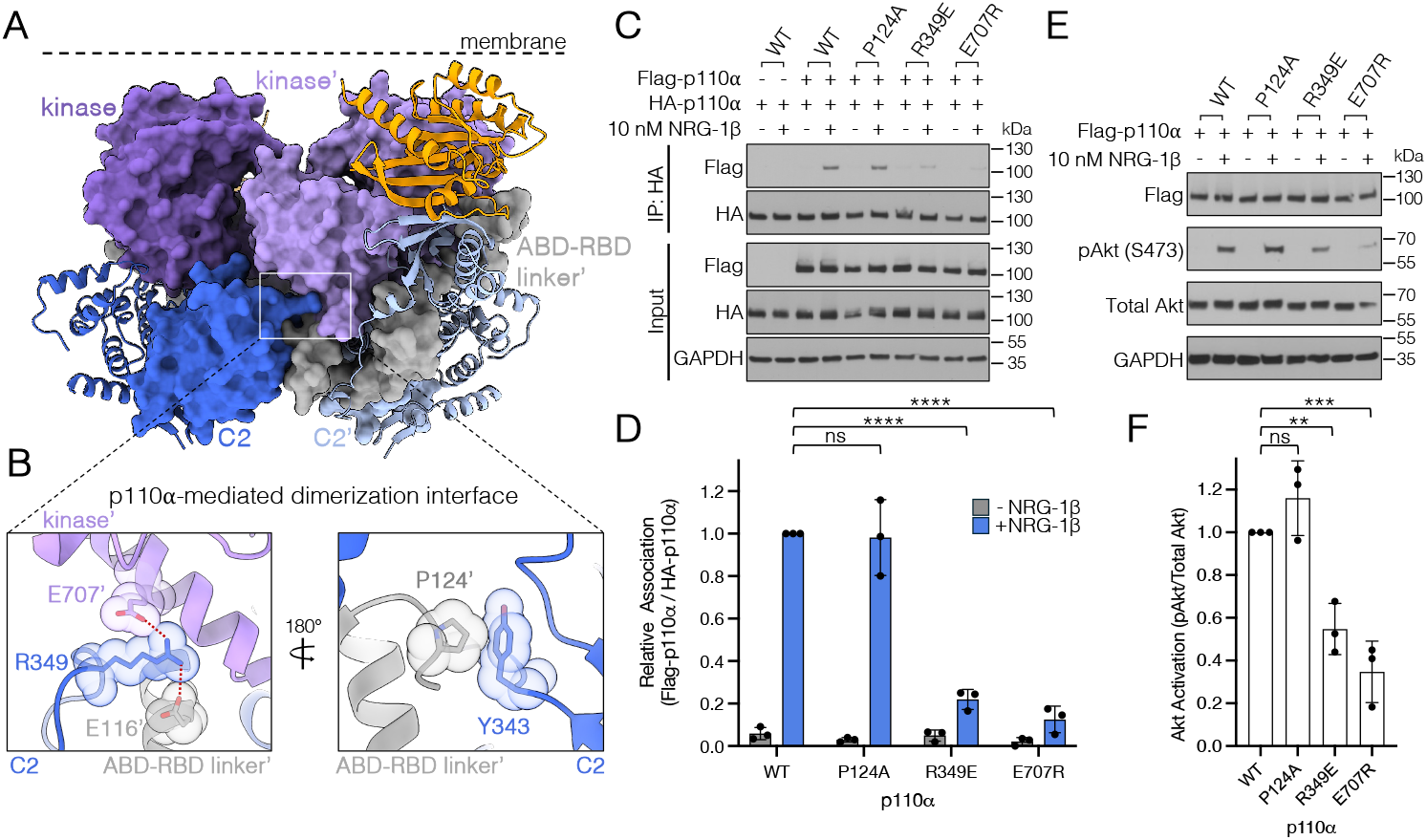
PI3Kα dimerization amplifies Akt signaling in cells. (**A**) The active, dimeric PI3Kα/KRas/phosphopeptide complex structure illustrating the ABD-RBD linker, C2 and kinase domains (shown in surface representation) which facilitate p110α- mediated dimerization. C2 domains are shown in light and dark blue, ABD-RBD linkers in light and dark gray, and kinase domains in light and dark purple. (**B**) Zoomed-in view of the dimerization interface, highlighting key interactions. (**C**) Co-immunoprecipitation of Flag-tagged and HA-tagged wild-type (WT) p110α and dimerization interface mutants of p110α (P124A, R349E, E707R) transiently expressed in MCF7 cells with and without NRG-1β stimulation and (**D**) corresponding quantification. (**E**) Akt phosphorylation induced by 10 nM NRG-1β stimulation in MCF7 cells transfected with Flag-tagged WT p110α or dimerization interface mutants of p110α (P124A, R349E, E707R) and (**F**) corresponding quantification. Protein levels (C, E) were detected with the indicated antibodies by Western Blot. Data in (D) and (F) are plotted as the mean with standard deviation from 3 independent experiments. Statistical significance was determined using One-way ANOVA Dunnett’s multiple comparisons test, **<0.01, ***p < 0.001, ****p < 0.0001.

Next, we investigated the contribution of PI3Kα homodimerization to signaling by measuring Akt phosphorylation in MCF7 cells co-expressing dimerization-deficient mutants, R349E and E707R, as a function of NRG1β stimulation. Akt activation was significantly impaired in cells expressing the dimerization-deficient PI3Kα mutants compared to cells expressing WT PI3Kα constructs upon NRG1β stimulation **(Fig. 4E, 4F)**. The P124A mutant, which did not disrupt PI3Kα dimerization **(Fig. 4C, 4D)**, did not affect Akt activation (**Fig. 4E, 4F**). Altogether, our results point to an important role of p110α-mediated PI3Kα homodimerization, via the interface revealed by our structure, in setting the amplitude of PI3Kα signaling.

## Discussion

Our nanodisc-bound PI3Kα/KRas structures outline discrete steps in the activation trajectory of PI3Kα and reveal how the PI3Kα holoenzyme uses its complex architecture to integrate multiple activating inputs (the membrane, Ras, PIP2 and the phosphopeptide) to achieve its fully activated state. In our structures, the disengagement of the inhibitory nSH2, iSH2, and ABD domains—often at quite a distance from the active site—is correlated with the precise re- organization of the active site. We show this more directly by performing focused 3D classification on each dataset with a mask encompassing just the active site of PI3Kα **(fig. S14)**. This analysis reveals that PI3Kα populates an equilibrium of states within each condition (POPC/POPS, POPC/POPS/PIP2, POPC/POPS/PIP2-phosphopeptide) **(fig. S15)** and that the conformation of helix kα12 is directly linked to the extent of the inhibitory interactions involving the nSH2, iSH2, and ABD domains—regions of the structure outside the mask used for classification. These findings suggest that differential engagement of p110α by p85α is sensed across the entire PI3Kα holoenzyme and communicated to the active site.

Our work brings new insights into the functional significance of helix kα12. Helix kα12 has been identified as a critical regulator of membrane binding and lipid kinase activity in class I and class III PI3Ks, but its precise mechanism of action has remained elusive (*25*,*33-35*). Interestingly, an inwardly rotated and helical helix kα12 is observed in the crystal structures of the autoinhibited class I PI3Ks, PI3Kβ and PI3Kγ **(fig. S8A)**. In contrast, in the structure of the class III PI3K, Vps34, helix kα12 is rotated away from the active site (*4*,*33*,*34*,*36*). This variability in the conformation of helix kα12 among PI3K isoforms, along with its involvement in crystal contacts with neighboring molecules, has clouded understanding of its position relative to the p110α active site and the membrane plane. Our structures show for the first time that the inward, helical conformation of helix kα12 constitutes a critical component of PI3Kα activation, stabilizing the PIP2 substrate and anchoring PI3Kα to the membrane.

The structure of the dimeric PI3Kα/KRas complex reveals a new role for the p110α subunit in regulating PI3Kα activation. PI3Kα dimerization has been reported before, though it was suggested to occur via the p85α subunit (*37*,*38*). p85α-driven p110α/p110β heterodimers were shown to preferentially bind PTEN and increase its activity, suggesting that such dimers negatively regulate the PI3K/AKT pathway (*39*,*40*). The p110α-mediated homodimer we report here corresponds to a fully activated PI3Kα state. We show that PI3Kα utilizes this newly described interface to dimerize in cells following growth factor stimulation and that PI3Kα/Akt signaling is weakened when this interface is broken. We posit that this p110α-mediated PI3Kα dimer forms only upon PI3Kα relocation to the membrane and represents a functionally distinct PI3Kα state that amplifies its signaling. This dimeric state might be particularly relevant for signaling downstream from multi-site PI3Kα scaffolds, like HER3 and Insulin Receptor Substrate (IRS), which have evolved to recruit multiple molecules of PI3Kα and, thus, have the capacity to capitalize on this regulatory mechanism.

Overall, this study shows that while Ras-mediated binding of PI3Kα to non-specific membranes partially disrupts p85α inhibitory contacts, the cSH2 domain is poised to block active site engagement of membrane lipids. This previously unknown role of the cSH2 domain in PI3Kα merits further investigation through higher resolution structural studies to determine its precise conformation. Binding of PIP2 induces a more activated state of PI3Kα; however, it remains susceptible to p85α rebinding and subsequent auto-inhibition, as evidenced by our 3D classification analysis **(fig. S15)**. It is only when PI3Kα is bound to phosphorylated YxxM motifs that the p85α binding interface is occluded by the dimeric p110α, preventing p85α rebinding and shifting the equilibrium fully toward the active state **(Fig. 5)**. Altogether, the newly defined conformational states of PI3Kα presented here not only offer new hypotheses for cell signaling, but also establish a much-needed foundation for designing novel therapeutics that target the unique features of both monomeric and dimeric active PI3Kα states.

**Fig. 5.**
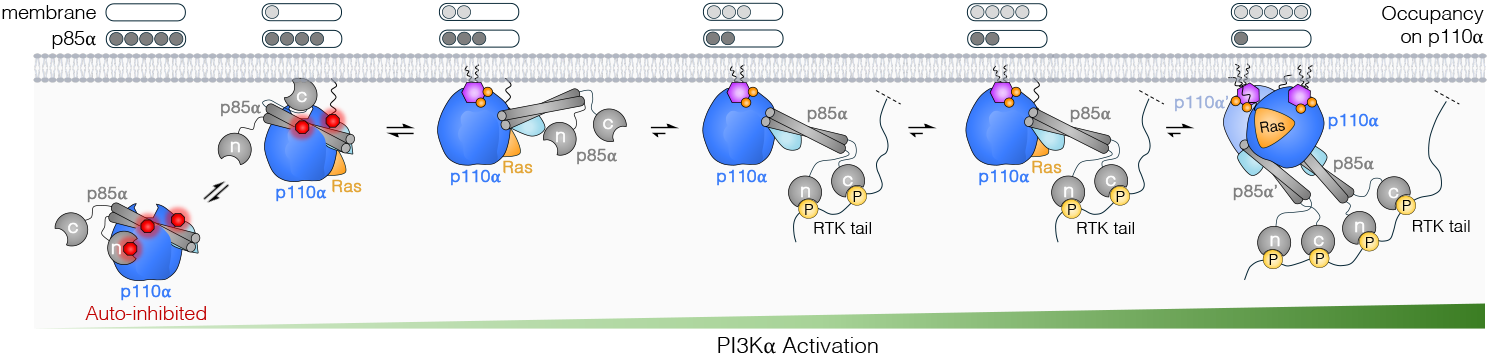
A model of PI3Kα activation at the membrane. Cartoon schematic depicting conformational states of PI3Kα along its activation trajectory and the relative occupancies of p85α and the membrane on p110α. p85α is shown in gray, p110α in blue, ABD of p110α in cyan, and Ras in orange. PIP2 is shown as a pink hexagon and inhibitory interfaces are depicted with red octagons.

## MATERIALS AND METHODS

### PI3Kα virus generation, expression, and purification

Full-length p85α and p110α were cloned into pFastBac1 and pFastBacHT B vectors, respectively. The p85α subunit is untagged while the p110α vector encodes an N-terminal 6X His tag. Baculovirus was generated for each construct separately using the DH10Bac system. The p85α and p110α heterodimer was co-expressed in Sf9 insect cells by co-infection of individual baculoviruses at 27ºC. Cells were harvested after 72 hrs, centrifuged, and flash frozen in liquid nitrogen for later use.

Pellets were resuspended in 50 mM Sodium Phosphate buffer, pH 8.0, 400 mM NaCl, 10 mM imidazole, 5% glycerol, 1% triton X-100, 10 mM βME. Cells were lysed by sonication at 40% amplitude for 5 min (10 sec ON, 5 sec OFF) and spun down in an Optima L-90K ultracentrifuge (Beckman Coulter) equipped with a 45 Ti rotor at 40,000 rpm for 1 hr at 4ºC. The clarified lysate was filtered using a 0.22 μm syringe filter and loaded onto a 1 ml HisTrap HP column equilibrated with binding buffer consisting of 50 mM Sodium Phosphate buffer pH 8.0, 400 mM NaCl, 10 mM Imidazole, 10 mM βME. The column was washed with 15 column volumes of binding buffer and eluted with binding buffer containing 250 mM Imidazole using a linear imidazole gradient over 35 column volumes. The eluate was diluted 2-fold with binding buffer without Imidazole and glycerol was added at a final concertation of 5%. The eluate was then concentrated using an Amicon Ultra-15 100k MWCO centrifugal filter and loaded onto a Superose 6 increase 10/300 GL column and separated using an isocratic gradient of 50 mM Tris pH 7.5, 150 mM NaCl, 10% glycerol, 1 mM TCEP. Peak fractions were concentrated using Amicon-Ultra 0.5 ml 100 kDa MWCO centrifugal filters.

### KRAS4b-FME virus generation, expression, purification, and GppNHp loading

A full-length farnesylated and methylated KRAS4b (KRAS-FME) was produced as previously described(*19*). Briefly, a clone for human KRAS4b (amino acids 2-188) with an upstream TEV protease cleavage site and an N-terminal 6XHis-MBP fusion tag was transformed into *E. coli* DE35 and plated on LB agar with 7μg/ml gentamicin, 50 μg/ml kanamycin, 10 μg/ml tetracycline, 100 μg/ml Bluo-gal, and 40 μg/ml IPTG. White colonies were grown in liquid culture, and bacmid DNA was purified by alkaline lysis. Positive clones were transfected into 100 ml of Tni-FNL insect cells at 1.5 x 10^6^ cells/ml, grown at 27°C for 4-6 days. The resulting P1 baculovirus was used to infect expression cultures, which were spun down and their pellets flash frozen after 72 hrs.

Pellets were resuspended in 20 mM HEPES, pH 7.3, 300 mM NaCl, 5 mM MgCl2, 1 mM TCEP, and 1 x protease inhibitor tablet (Sigma Aldrich). Cells were lysed using an EmulsiFlex-C5 homogenizer (Avestin), and the cell lysate was clarified by ultracentrifugation at 50,000 xg for 30 min, at 4 °C. Imidazole was added to the supernatant at a final concertation of 35 mM and loaded onto a 10 mM IMAC column (HisTrap, GE Healthcare) equilibrated with 20 mM HEPES, pH 7.3, 300 mM NaCl, 5 mM MgCl_2_, 1 mM TCEP, 35 mM imidazole, and 1x protease inhibitor tablet. The column was washed with equilibration buffer and proteins eluted with increasing imidazole gradient up to 500 mM imidazole. Peak fractions were dialyzed into 20 mM MES, pH 6.0, 150 mM NaCl, 5 mM MgCl_2_, and 1 mM TCEP and diluted into dialysis buffer to reduce salt concentration to 75 mM NaCl. Dialyzed protein was loaded onto a 10 ml SP Sepharose column equilibrated with 20 mM MES, pH 6.0, 75 mM NaCl, 5 mM MgCl_2_, and 1 mM TCEP. The column was washed and then eluted with a salt gradient of up to 650 mM NaCl. Protein was incubated with a 1:10 molar ratio of TEV protease and simultaneously dialyzed into 20 mM HEPES, pH 7.3, 300 mM NaCl, 5 mM MgCl_2_, and 1 mM TCEP overnight at 4°C. Digested protein was separated by IMAC, peak fractions pooled, concentrated using an Amicon-Ultra10 kDa MWCO centrifugal filters, and flash frozen for storage at -80°C.

Purified KRas was diluted in HEPES buffer containing 200 mM ammonium sulfate and 100 μM ZnCl_2_ until the MgCl_2_ concentration was below 1 mM. The diluted protein was then mixed with a 10:1 molar ratio of GppNHP to KRas and incubated with alkaline phosphatase agarose beads (1 unit per mg of KRas) at room temperature for 3 hrs. Beads were spun down at 2000xg for 2 min and the supernatant was incubated with 10-fold molar excess of GppNHp in the presence of 5 mM MgCl_2_ overnight at 4°C . Excess GppNHp was removed with a PD-10 G-25 desalting column (Cytiva, 17085101) equilibrated with 20mM HEPES, 300mM NaCl, 1mM MgCl_2_, 1mM TCEP. The nucleotide exchange efficiency was determined by high-performance liquid chromatography.

### Nanodisc preparation

Chloroform/methanol solutions of POPC, POPS, and PIP2 (Avanti Polar Lipids: 850457, 840034, 840046) were prepared and mixed together in the appropriate molar ratios (65:35 POPC:POPS for non-PIP2 nanodiscs and 70:25:5 POPC:POPS:PIP2 for PIP2 nanodiscs) in glass vials. Lipids were initially dried down under nitrogen gas and left overnight in a freeze dryer for complete solvent evaporation. Lipid films were resuspended in 20 mM Tris, pH 7.5, 100 mM NaCl, 0.5 mM EDTA, 40 mM Sodium Cholate to a final lipid concentration of 20 mM. Resuspended lipids were vortexed and sonicated in a bath sonicator for 45 min for solubilization. Detergent-solubilized lipids were incubated with MSP1E3D1 protein (Sigma-Aldrich, M7074) at a 1:130 (non-PIP2 lipid mixtures) and 1:140 (PIP2-conatining lipid mixtures) molar ratio (MSP:lipid) for 1 hr on ice. Bio-beads (Bio-Rad, 1528920) were activated in methanol for 30 min while shaking and thoroughly washed with ddH2O and 20 mM Tris, pH 7.5, 100 mM NaCl. Bio-beads were then added to the lipids-MSP mixtures at a final concentration of 0.8 mg/ml and incubated overnight at 4 °C with shaking. Nanodiscs were purified using either a Superdex 200 increase 10/300 GL or Superose 6 increase 10/300 GL column with an isocratic gradient of 50 mM Tris pH 7.5, 150 mM NaCl, 1 mM TCEP. Peak fractions were concentrated using Amicon-Ultra 0.5 ml 30 kDa MWCO centrifugal filters.

### Complex formation and sample preparation for cryo-electron microscopy imaging

PI3Kα, KRAS4b-FME and molecular glue were incubated at final concentrations of 3 μM, 6 μM, and 30 μM, respectively, for 1 hr on ice. Nanodiscs were added to the protein complex mixture to a final concentration of 6 μM and incubated for an additional 45 min at RT. PI3Kα- KRAS4b-FME-nanodisc complexes were loaded onto a Superdex 200 Increase 3.2/300 GL column and separated using an isocratic gradient of 50 mM Tris pH 7.5, 150 mM NaCl, 1 mM TCEP. For bis-phosphopeptide-bound complexes, PI3Kα, KRAS4b-FME, and molecular glue were incubated with a HER3-derived bis-phosphopeptide (GD(p)YAAMGACPASEQG(p)YEEMRA, Genscript) at a final concentration of 15 μM for 1 hr on ice, followed by incubation with nanodiscs and separation by size exclusion chromatography. The peptide-bound complex was also prepared by forming PI3Kα-KRAS4b-FME-nanodisc complex first, followed by incubation with bis-phosphopeptide for 1 hr on ice and by separation by size exclusion chromatography. The order of peptide addition did not affect the resulting PI3Kα-KRAS4b-FME-nanodisc complex cryo-EM maps.

Peak fractions of each complex were pooled and diluted to a 40 mAU equivalent concentration. Immediately before preparing cryo-EM grids, fluorinated octyl maltoside (Anatrace, O310F) was added to the samples at a final concentration of 0.0125% to alleviate orientation bias of nanodisc-bound complexes. For the POPC/POPS nanodisc complex, 3 μl of sample was applied to graphene-oxide coated Quantifoil R1.2/1.3 300 mesh Au holey-carbon grids, blotted using a Vitrobot Mark IV (FEI), and plunge frozen in liquid ethane at 5 °C, 100% humidity, 30 sec wait time, 7 sec blot time, and 0 blot force. For the POPC/POPS/PIP2 nanodisc complexes, in the presence or absence of bis-phosphopeptide, 3 μl of sample was applied to graphene-oxide coated Quantifoil R1.2/1.3 300 mesh Au holey-carbon grids functionalized with PEG-amine, to combat the excess negative charge from PIP2 and bis-phosphopeptide. Grids were blotted and plunge frozen in liquid ethane at 5 °C, 100% humidity, 30 sec wait time, 3 sec blot time, and 0 blot force.

Grids were imaged on a 300-keV Titan Krios (FEI) with a K3 direct electron detector (Gatan) and a BioQuantum energy filter (Gatan) using SerialEM v4.1 and Digital Micrograph v3.2. Data for the PI3Kα-KRAS4b-FME-nanodsic complexes were collected in non-super-resolution mode at a physical pixel size of 0.8189 Å pixel^−1^, with a dose rate of 16.0 e^−^ pixel^−1^ s^−1^ (operated in non-CDS mode), at a nominal magnification of 105,000x. Images were recorded with a 2 sec exposure over 80 frames with a dose rate of 0.57 e^−^ Å^−2^ per frame.

### Image processing and 3D reconstruction

Raw movies were corrected for motion and radiation damage with MotionCor2(*41*) during collection launched via Scipion(*42*) (developer version), and the resulting sums were imported into CryoSPARC v4.3.1. Micrographs were curated based on ice thickness, CTF fit resolution, and particle distribution. Micrograph contrast transfer function (CTF) parameters were estimated with the patch CTF job in CryoSPARC. The following is a brief description of the general processing workflow for all datasets. See processing workflow charts for further details **(Figs. S1, S6, and S10)**. Particles were initially picked with blob picking, extracted, and 2D-classified yielding templates for template picking. Particles were then template picked with low-pass filtered (to 20 Å) 2D class averages and resulting picks were extracted with 4x Fourier cropping and subjected to iterative rounds of *ab initio* and heterogeneous refinements. Refined particle stacks were then used to train Topaz v0.2.5 and extract 4x binned particles from Topaz-based particle picking. These particles were subjected to iterative rounds of *ab initio*, heterogeneous refinements and 3D classification. Finally, unbinned particles were re-extracted and run through non-uniform refinements to achieve reconstruction with the highest resolution. The final reconstructions of the POPC/POPS-bound PI3Kα/KRas class 1, class 2, and class 3 complexes used for model building included 111,772, 114,960, and 117,424 particles with C1 symmetry and resulted in overall resolutions of 3.01 Å, 3.23 Å, and 3.05 Å by gold standard FSC (GS-FSC) cut-off of 0.143 (masked), sharpened with -103.5, -111.9, and -103.3 B-factors, respectively. The final reconstruction of the POPC/POPS/PIP2-bound PI3Kα/KRas complex used for model building included 223,376 particles with C1 symmetry and resulted in an overall resolution of 2.82 Å, sharpened with -108.3 B-factor. The POPC/POPS/PIP2-bound PI3Kα/KRas/peptide monomer and dimer complexes used for model building included 111,698 and 280,864 particles with C1 and C2 symmetry, respectively, and resulted in overall resolutions of 2.89 Å and 2.61 Å, sharpened with - 101.9 and -101.5 B-factors. Finally, each map was assessed for local and directional resolutions through ResMap v1.1.461 and 3DFSC v3.062 server, respectively.

### Model building, refinement, and validation

Model building for PI3Kα/KRas complexes began with the alignment of the AlphaFold2 models of p85α, p110α, and KRas onto the p110γ/HRas crystal structure (1HE8). The resulting models were then rigid body fit into the corresponding 3D volumes in ChimeraX 1.7.1. The models were then flexibly fit into their respective cryo-EM maps with FastRelax protocol in Rosetta 3.0(*43*) in torsion space and then were further manually examined and refined in Coot and ISOLDE(*44*). Per-atom B-factors were assigned in Rosetta, indicating the local quality of the map around that atom. The residues modeled for the high resolution models of POPC/POPS-bound complexes are as follows: Class 1 (chain A: 8-312, 323-412, 416-502, 522-864, 873-1056; chain B: 446-586; chain C: 2-172), Class 2 (chain A: 19-56, 60-72, 80-83, 94-311, 323-411, 417-502, 522-862, 873-940, 951-1049; chain B: 481-507, 520-545, 560-568; chain C: 2-172), Class 3 (chain A: 107-311, 323-411, 416-501, 523-864, 873-942, 950-1050; chain C: 2-172). The residues modeled for the high-resolution models of the POPC/POPS/PIP2-bound complex are as follows: chain A (107-310, 323-502, 523-940, 952-1067), chain C (2-172). The residues modeled for the high-resolution models of the POPC/POPS/PIP2/peptide-bound complexes are as follows: monomer (chain A: 108-310, 323-347, 351-411, 417-502, 523-940, 951-1067; chain C: 2-172), dimer (chain A: 108-310, 323-411, 416-502, 519-941, 952-1067; chain B: 108-311, 323-411, 416- 502, 519-941, 952-1067, chain C: 1-172, chain D: 2-172). MolProbity, as part of the Phenix 1.19.2 validation tools, was used to assess the quality of the models. Q-scores were calculated using the appropriate plugin in ChimeraX 1.7.1.

Model building for the low resolution POPC/POPS-bound class 1 PI3Kα/KRas complex began with AlphaFold3 models of p85α, p110α, and KRas, which were rigid body fit into a 10 Å low-pass filtered class 1 volume in ChimeraX. The SH3 and BH domains of p85α (residues 1-323) were deleted from the model as they were located outside of the map and on the opposite side of the extra density observed between the kinase domain and the nanodisc. The model was energy-minimized in Rosetta without cryo-EM map first before being flexibly fit in torsion space into the 10 Å cryo-EM map. The nSH2 and cSH2 domains from the resulting model were then combined with the high resolution POPC/POPS-bound class 1 model.

To model the activation loop in the PIP2-bound complex, p110α residues 941-950 were first manually built in Coot as glycines into a 5 Å low-pass filtered map. 7000 models of the loop were generated using the iterative rebuild protocol in Rosetta and filtered by Rosetta energy and density fit scores. The top 20 models were inspected manually and the best fitting loop was mutated back to p110α residues and was further refined in ISOLDE. The final loop was combined with the high-resolution model of the PIP2-bound complex. To model the activation loop in the peptide-bound monomeric and dimeric complexes, the PIP2-bound complex and peptide-bound complexes were aligned in ChimeraX, and the PIP2-bound loop was stitched onto the high-resolution models of the peptide-bound complexes. The peptide-bound complexes were then flexibly fit into their respective 5 Å low-pass filtered cryo-EM maps with FastRelax protocol in torsion space in Rosetta followed by further refinement in Coot and ISOLDE.

### Analysis of conformational variability of helix kα12 in existing PI3K structures

Using the “Similar Structures” tool in ChimeraX 1.9, and the p110α subunit from our PIP2- bound model as a template, all structures in the PDB that had a BLAST alignment score better than the 1e-200 cutoff were downloaded, without trimming chain ends. The structures were then clustered based on the UMAP plot with the following command: “similarstructures cluster #1/A:1032-1044, 975-993, 891-910 clusterDistance 1 clusterCount 4” to specifically cluster based on conformations of helix kα11, helix kα10 and helix kα8 **(Fig. S8A)**. Similar workflow was used for **Figure S8B** but focusing just on PI3Kα structures and using 3 clusters.

### Plasmids and cell culture

The human 3xFlag-PIK3CA, Myc-PIK3R1, and HA-PIK3CA cDNAs were synthesized by GenScript and Sino Biological, and subcloned into pcDNA3.1 and pCMV3 vectors, respectively. Mutations were introduced using Quikchange mutagenesis (Agilent). All constructs were verified via whole plasmid DNA sequencing (Plasmidsaurus). MCF7 cells were maintained at 37 ºC and 5% CO_2_ and cultured in Dulbecco’s modified Eagle media (Life Technologies) supplemented with 10% FBS (Hyclone) and 1% penicillin/streptomycin (Life Technologies). PIK3CA and PIK3R1 constructs were transiently transfected into cells 24 hrs prior to cell lysis using Lipofectamine 3000 (Invitrogen; co-immunoprecipitation assays) or Lipofectamine LTX (Invitrogen; signaling assays) according to the manufacturer’s protocol.

### Co-immunoprecipitation and Western blot analysis

MCF7 cells were seeded in 100 mm dishes such that they were at 70% confluency for transfection the following day. 24 hrs post-transfection, cells were serum starved with 0.1 % FBS for 16 hrs. Following starvation, cells were stimulated with 10 nM NRG-1β (Peprotech, 100- 03) for 10 min. Cells were washed two times on ice with ice-cold PBS followed by lysis (50 mM Tris pH 8.0, 150 mM NaCl, 0.5% Triton X-100, 0.5% NP-40, 1 mM NaF, 1 mM Na_3_VO_4_, 1 mM EDTA, cOmplete mini EDTA-free protease inhibitor cocktail [Roche]). Cells were scraped and incubated with rotation at 4 °C for 45 min to ensure complete lysis. Lysates were clarified by centrifugation for 10 min at 15,000 rpm at 4 °C. The clarified lysates were pre-cleared with 100 μL Protein A beads (Novex) for 30 min at 4 °C with rotation. The pre-cleared lysates were then incubated with 100 μL of antibody/protein A complexed beads (anti-HA [mouse, SCBT, sc- 7392]) per 100 mm plate overnight at 4 °C with rotation. The beads were washed three times with lysis buffer. The bound proteins were eluted from the beads using SDS-loading buffer and were boiled at 95 °C for 10 min prior to SDS-PAGE and analysis by Western blot. Samples were run on 4-15% acrylamide gels and transferred onto PVDF membranes. Membranes were blocked in 5% milk in TBS and 0. 1% Tween-20 (TBS-T) for 30 min at room temperature, followed by incubation with primary antibodies diluted in blocking buffer (anti-HA [rabbit, CST, 3724S], anti-GAPDH [rabbit, CST, 2118S]) at 4 °C overnight. Next, blots were incubated with secondary antibodies diluted in blocking buffer (anti-Flag HRP [Sigma Aldrich, A8592], anti-rabbit [CST, 7074S]) for 1 hr at room temperature. ECL Western blotting detection reagent (Cytiva) or ECL prime (Cytiva) were used for detection.

Western blot quantification was performed in FIJI Version 2.0.0 using the gel analysis tool. First, the Flag IP signal was divided by the corresponding HA IP signal. Next, each Flag IP/HA IP signal was normalized against the WT Flag IP/HA IP signal. Statistical significance was determined using One-way ANOVA Dunnett’s multiple comparisons test in Prism 9 (Graphpad Software, Inc.).

### AKT signaling and Western blot analysis

MCF7 cells were seeded in 6-well dishes such that they were at 70% confluency for transfection the following day. 24 hrs post transfection, cells were serum starved with 0.1 % FBS for 16 hrs. Following starvation, cells were stimulated with 10 nM NRG-1β for 10 min. Cells were washed two times on ice with ice cold PBS followed by lysis (50 mM Tris pH 8.0, 150 mM NaCl, 1% NP-40, 0.5% sodium deoxycholate, 0.1% SDS, 1 mM NaF, 1 mM Na_3_VO_4_, 1 mM EDTA, cOmplete mini EDTA-free protease inhibitor cocktail [Roche]). Cells were scraped and lysed on ice for 10 min. Lysates were clarified by centrifugation for 10 min at 15,000 rpm at 4 °C. Clarified lysates were boiled for 10 min with SDS loading buffer prior to SDS-PAGE and analysis by Western blot. Samples were run on 4-15% acrylamide gels and transferred onto PVDF membranes. Membranes were blocked in 5% milk in TBS and 0. 1% Tween-20 (TBS-T) for 30 min at room temperature, followed by incubation with primary antibodies diluted in blocking buffer (anti-pan Akt [mouse, CST, 2920S]; anti-phospho-Akt, Ser 473 [rabbit, CST, 9271S], anti-GAPDH [rabbit, CST, 2118S]) at 4 °C overnight. Next, blots were incubated with secondary antibodies diluted in blocking buffer (anti-Flag HRP [Sigma Aldrich, A8592], anti-rabbit HRP [CST, 7074S], anti-mouse HRP [CST, 7076S]) for 1 hr at room temperature. ECL Western blotting detection reagent (Cytiva) or ECL prime (Cytiva) were used for detection.

Western blot quantification was performed in FIJI Version 2.0.0 using the gel analysis tool. First, the phospho-Akt signal was divided by the corresponding Total Akt signal. Next, each phospho-Akt/Total Akt signal was normalized against the WT phospho-Akt/Total Akt signal. Statistical significance was determined using One-way ANOVA Dunnett’s multiple comparisons test in Prism 9 (Graphpad Software, Inc.).

## Acknowledgments

We thank the NCI Ras initiative (Dwight Nissley, Dominic Esposito, and Andy Stephen) for initially providing KRas-FMe and for subsequent training, protocols, and reagents enabling in-house purification of KRas-FMe; Frank McCormick for providing the glue compound, helpful discussions, and critical reading of the manuscript; Komal Pawar for providing technical advice on complex reconstitution; the Jura and Verba labs for helpful discussions, and the UCSF Cryo-EM core (David Bulkley, Glenn Gilbert, Li Wang, and Matt Harrington) for help in collecting data.

## Funding

NIH U54CA274502 to NJ and QBI funding to KAV

## Author contributions

Conceptualization: HT, NJ, KAV

Methodology: HT, NJ, KAV

Investigation: HT, MDP, AM, BGM

Visualization: HT, NJ, KAV

Funding acquisition: NJ, KAV

Project administration: NJ, KAV

Supervision: NJ, KAV

Writing – original draft: HT, NJ, KAV

Writing – review & editing: HT, MDP, AM, BGM, NJ, KAV

## Competing interests

Authors declare they have no competing interests.

## Data and materials availability

The data that supports this study is available from the corresponding authors upon reasonable request. Cryo-EM maps have been deposited in the Electron Microscopy Data Bank (EMDB) under accession codes EMD-49454 (POPC/POPS Class 1), EMD-49456 (POPC/POPS Class 2), EMD-49455 (POPC/POPS Class 3), EMD-49451 (PIP2), EMD-49453 (PIP2/peptide monomer), EMD-49452 (PIP2/peptide dimer), EMD-49519 (10 Å POPC/POPS Class 1), EMD-49460 (5 Å PIP2), EMD-49459 (5 Å PIP2/peptide monomer), EMD-49458 (5 Å PIP2/peptide dimer). Associated models have been deposited in the Protein Data Bank (PDB) with accession codes 9NI6 (POPC/POPS Class 1), 9NI8 (POPC/POPS Class 2), 9NI7 (POPC/POPS Class 3), 9NI3 (PIP2), 9NI5 (PIP2/peptide monomer), 9NI4 (PIP2/peptide dimer), 9NLC (10 Å POPC/POPS Class 1), 9NIF (5 Å PIP2), 9NIE (5 Å PIP2/peptide monomer), 9NID (5 Å PIP2/peptide dimer).

## List of Supplementary Materials

Figs. S1 to S15

Tables S1 to S3

## Notes

### Competing Interest Statement

The authors have declared no competing interest.

