## Supplemental Materials for "Structures of the PI3Kα/KRas complex on lipid bilayers reveal the molecular mechanism of PI3Kα activation"

**The PDF file includes:**

Figs. S1 to S15
Tables S1 to S3

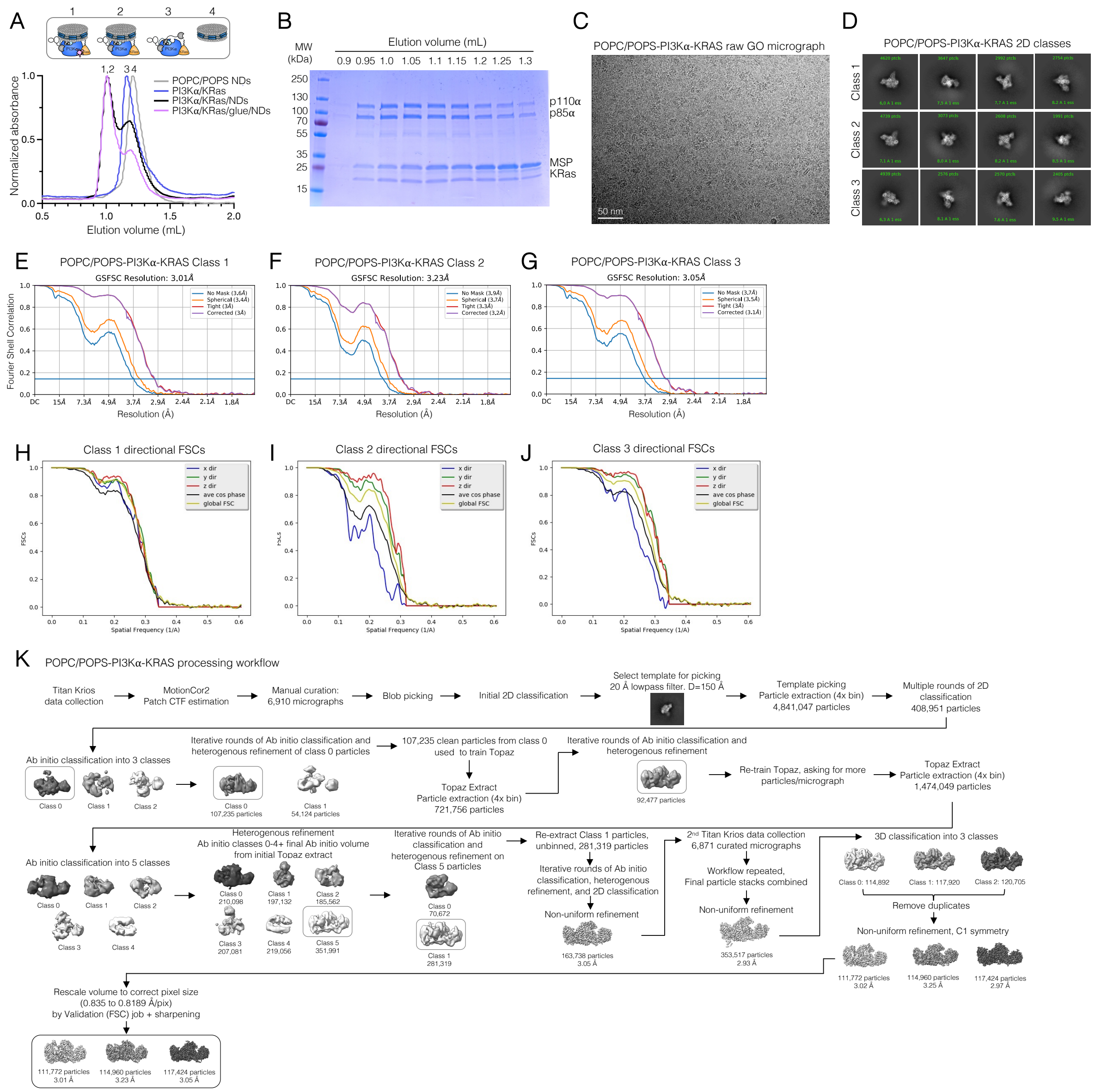


**Fig. S1. In-vitro reconstitution, map quality and processing workflow for the POPC/POPS-PI3Kα/KRas complex dataset.** (**A**) Representative size exclusion chromatography profiles of POPC/POPS/MSP1E3D1 nanodiscs (gray), PI3Kα/KRas complex (blue), PI3Kα/KRas complex on nanodiscs (black), and PI3Kα/KRas/molecular glue complex on nanodiscs (pink) resolved on a Superdex 200 Increase 3.2/300 column. (**B**) Corresponding Coomassie-stained SDS-PAGE analysis of the purified PI3Kα/KRas/molecular glue complex on nanodiscs. (**C**) Representative micrograph of the POPC/POPS-PI3Kα/KRas/glue complex sample on Quantifoil R1.2/1.3 300 mesh Au holey-carbon graphene oxide grids from a dataset of 13,781 micrographs. The scale bar corresponds to 50 nm. (**D**) Representative cryo-EM 2D class averages of particles corresponding to class 1, class 2, and class 3 complexes. (**E-G**) Gold Standard Fourier Shell Correlation (GSFSC) of final maps used for model building for class 1 (E), class 2 (F), and class 3 (G) complexes from CryoSPARC 4. (**H-J**) Directional FSCs of class 1 (H), class 2 (I) and class 3 (J) complexes calculated by the 3DFSC server. (**K**) Processing workflow of POPC/POPS-PI3Kα/KRas/glue complex dataset.

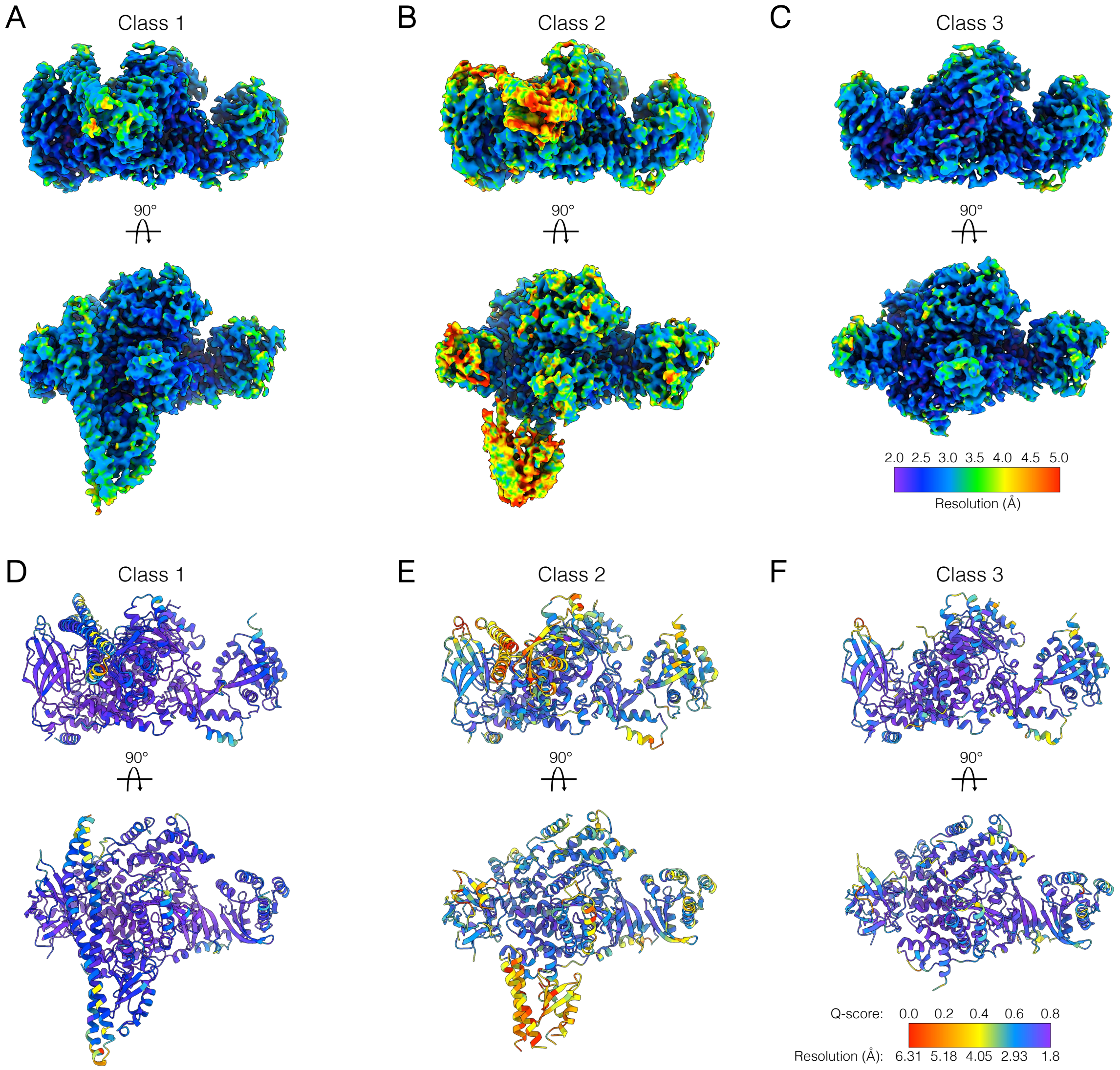


**Fig. S2. Resolution estimation and Q-score analysis of the POPC/POPS-PI3Kα/KRas complexes.** (**A-C**) Cryo-EM maps of the class 1 (A), class 2 (B), and class 3 (C) complexes colored according to local resolution determined by ResMap. (**D-F**) Corresponding models of class 1 (D), class 2 (E), and class 3 (F) complexes colored by estimated per residue Q-score. Color bars indicate corresponding resolution in Å (A-C) or estimated resolution in Å for reported Q-scores (D-F).

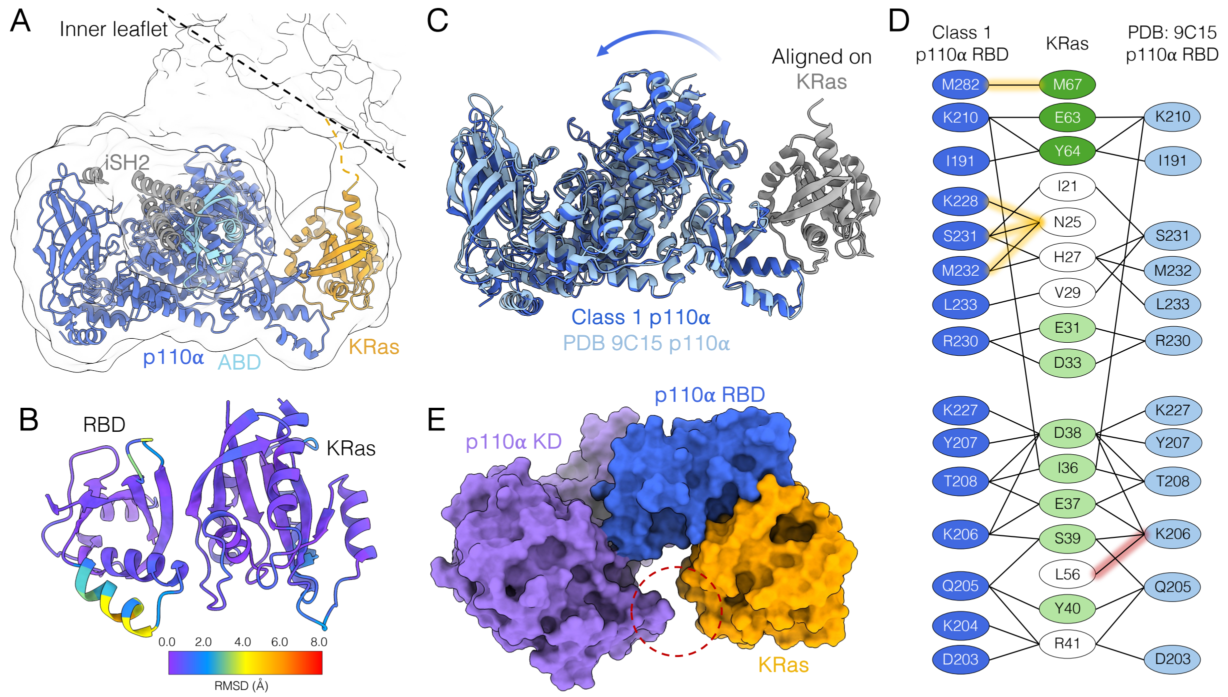


**Fig. S3. Comparison of the PI3Kα/KRas interface with and without a lipid bilayer.** (**A**) The structure of the class 2 PI3Kα/KRas complex overlayed with its 10 Å low-pass filtered cryo-EM map, highlighting insertion of KRas C-terminus into the membrane. p110α is shown in blue and its ABD in cyan, iSH2 domain of p85α in gray, and KRas in orange. (**B**) The POPC/POPS-bound class 1 structure RBD/KRas region colored by per residue Root Mean Square Deviation (RMSD, Å) relative to the RBD/KRas region of the p110α/KRas solution crystal structure (PDB ID: 9C15), demonstrating the rigidity of the RBD/KRas interface. (**C**) Overlay of the POPC/POPS-bound class 1 structure and p110α/KRas solution crystal structure (9C15) aligned on KRas, showcasing the rotation of the p110α catalytic core away from the RBD/KRas interface upon membrane binding. KRas is colored in gray, the class 1 p110α in blue, and 9C15 p110α in light blue. (**D**) Interactions comprising the PI3Kα/KRas interface in the POPC/POPS-bound class 1 structure and p110α/KRas solution crystal structure (9C15). KRas switch I residues are in light green and switch II residues in dark green. Unique contacts are highlighted in orange and red for the class 1 and crystal structures, respectively. (**E**) Class 1 PI3Kα/KRas structure shown in surface representation. The p110α RBD is shown in blue, kinase domain in purple, and KRas in orange. The remaining domains of the complex are omitted for clarity. The absence of KRas/kinase domain contacts is highlighted with a red circle.

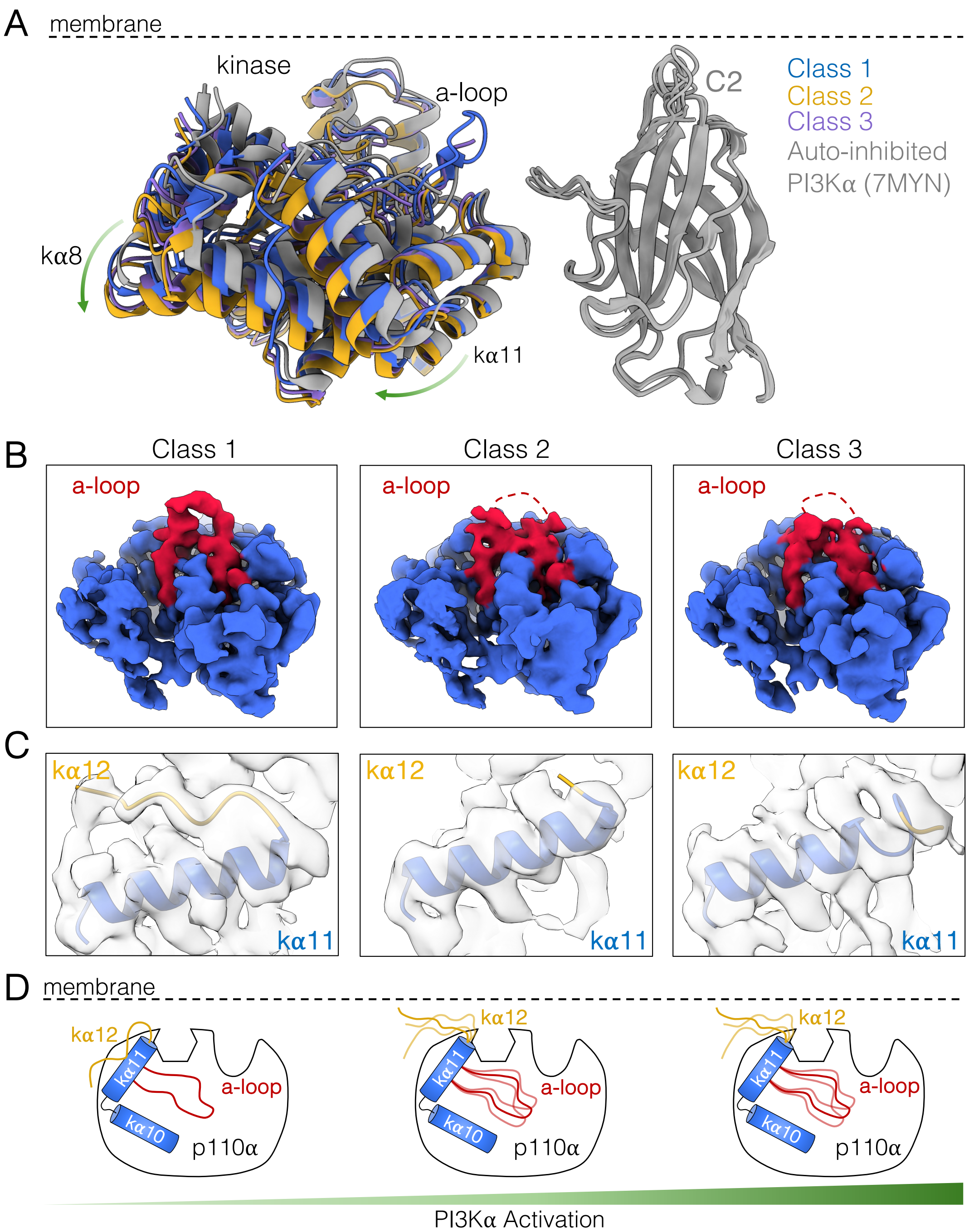


**Fig. S4. Conformational rearrangements in PI3Kα upon binding to KRas and POPC/POPS nanodiscs.** (**A**) Overlay of POPC/POPS-bound structures with the full-length, solution cryo-EM structure of PI3Kα (PDB ID: 7MYN). Structures are aligned on the C2 domain. 7MYN is shown in gray, class 1, class 2, and class 3 kinase domains in blue, yellow, and purple, respectively. The remaining PI3Kα domains and KRas are not shown for clarity. (**B-C**) 4 Å lowpass filtered cryo-EM maps of class 1, class 2, and class 3 reconstructions overlayed with their corresponding models showing an increasingly dynamic activation loop (B; shown in red) and helix kα12 (C; shown in yellow). (**D**) Cartoon schematic depicting conformational changes within the active site of PI3Kα along the three POPC/POPS-bound classes with p85α omitted for clarity.

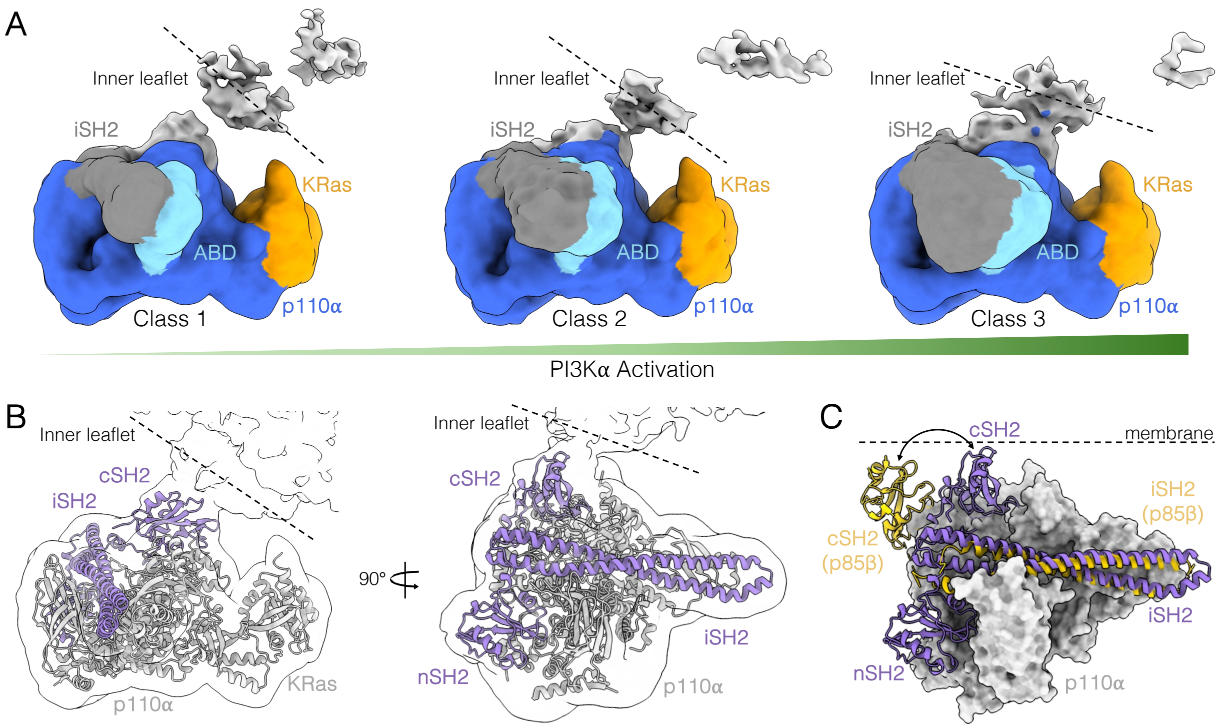


**Fig. S5. PI3Kα/nanodisc interactions in POPC/POPS-bound classes.** (**A**) 10 Å low-pass filtered maps of class 1, 2 and 3 POPC/POPS-bound complexes illustrating orientation of the nanodisc inner leaflet relative to the PI3Kα active site across the three classes. p110α is colored in blue, its ABD domain in cyan, p85α-iSH2 in gray, and KRas in orange. (**B**) Structure of the full-length class 1 PI3Kα/KRas complex overlayed with its 10 Å low-pass filtered cryo-EM map, illustrating placement of the p85α cSH2 domain within the extra density between the p110α active site and the nanodisc inner leaflet. (**C**) Structure of the class 1 PI3Kα/KRas complex overlayed with the solution crystal structure of icSH2 PI3Kβ, PDB ID: 2Y3A (aligned on p110α), illustrating the rotation of the p85α cSH2 domain toward the membrane. In (B-C), p85α is colored in purple, p110α and KRas in gray, and p85β in yellow.

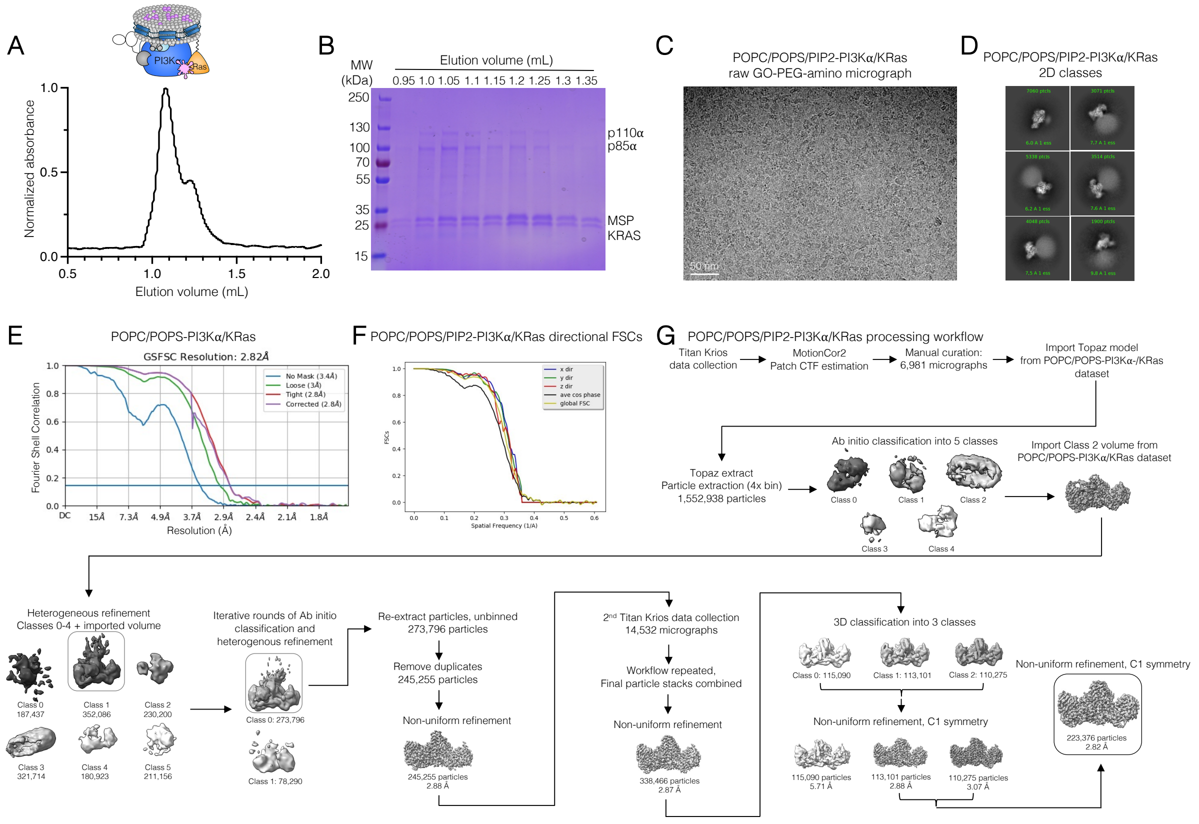


**Fig. S6. In-vitro reconstitution, map quality and processing workflow for the POPC/POPS/PIP2-PI3Kα/KRas complex dataset.** (**A**) Representative size exclusion chromatography profile of PI3Kα/KRas/molecular glue complex on POPC/POPS/PIP2/MSP1E3D1 nanodiscs resolved on a Superdex 200 Increase 3.2/300 column. (**B**) Corresponding Coomassie-stained SDS-PAGE analysis of the purified PI3Kα/KRas/molecular glue complex. (**C**) Representative micrograph of the POPC/POPS-bound PI3Kα/KRas complex sample on Quantifoil R1.2/1.3 300 mesh Au holey-carbon PEG-amine functionalized graphene oxide grids from a dataset of 21,513 micrographs. The scale bar corresponds to 50 nm. (**D**) Representative cryo-EM 2D class averages of particles corresponding to the POPC/POPS/PIP2-PI3Kα/KRas complex. (**E**) Gold Standard Fourier Shell Correlation (GSFSC) of final map used for model building for the POPC/POPS/PIP2-PI3Kα/KRas complex from CryoSPARC 4. (**F**) Directional FSCs of the POPC/POPS/PIP2-PI3Kα/KRas complex calculated by the 3DFSC server. (**G**) Processing workflow of POPC/POPS/PIP2-PI3Kα/KRas complex dataset.

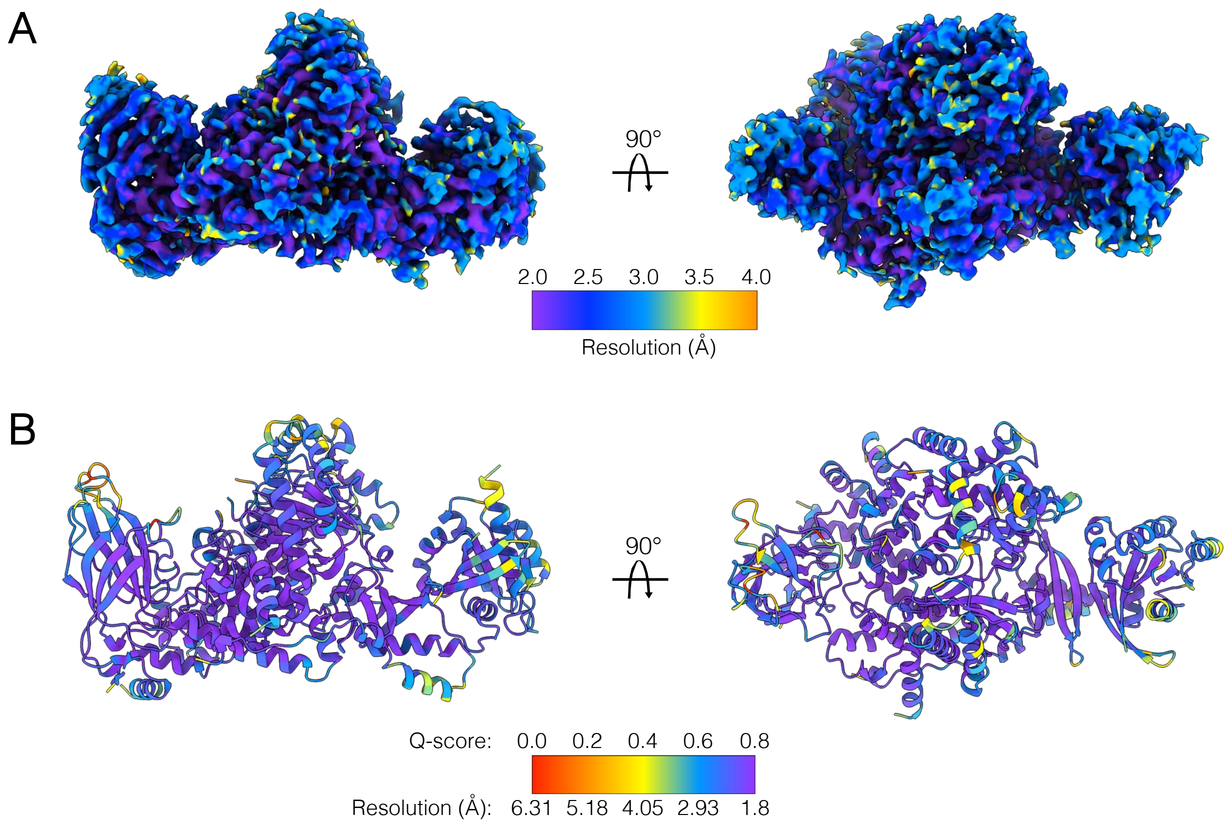


**Fig. S7. Resolution estimation and Q-score analysis of the POPC/POPS/PIP2-PI3Kα/KRas complex.** (**A**) Cryo-EM map of the POPC/POPS/PIP2-bound PI3Kα/KRas complex colored according to local resolution determined by ResMap. (**B**) Corresponding model of the POPC/POPS/PIP2-bound PI3Kα/KRas complex colored by estimated per residue Q-score. Color bars indicate corresponding resolution in Å (A) or estimated resolution in Å for reported Q-scores (B).

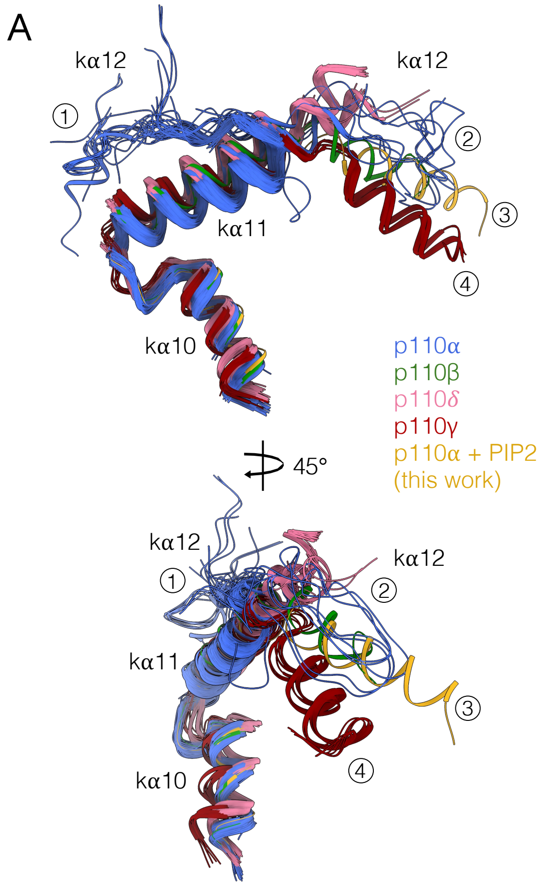


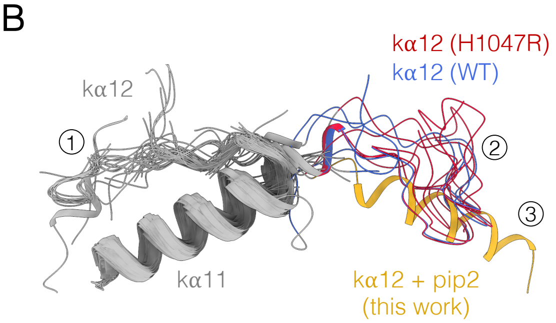


C

| PDB ID | p110α | p85α | Inhibitor? |
|---|---|---|---|
| 3HIZ | H1047R | niSH2 | no |
| 3HHM | H1047R | niSH2 | yes |
| 8V8U | H1047R | niSH2 | yes |
| 8V8J | H1047R | niSH2 | yes |
| 8V8I | H1047R | niSH2 | yes |
| 8V8H | H1047R | niSH2 | yes |
| 8TGD | H1047R | niSH2 | yes |
| 4A55 | WT | iSH2 | yes |
| 8TDU | WT | niSH2 | yes |
| 5DXH | WT | iSH2 | yes |

**Fig. S8. Conformational variability of helix kα12 among Class I PI3Ks.** (**A**) Structures of Class I PI3Ks aligned on p110α helices kα8 and kα11. Four main clusters of helix kα12 conformations are highlighted: disordered/outward (cluster 1), disordered/inward (cluster 2), helical/inward (cluster 3), and helical/acutely inward (cluster 4). Published p110α structures are shown in blue, p110β in green, p110δ in pink, p110γ in red, and the PIP2-bound p110α structure from this work in yellow. (**B**) Published structures of PI3Kα aligned on p110α helices kα8 and kα11, highlighting the presence of clusters 1-3. Cluster 1 structures are shown in gray, cluster 2 in blue (WT) and red (H1047R), and the PIP2-bound p110α from this work in yellow. (**C**) Cluster 2 published stuctures of PI3Kα, highighting the predominence of the H1047R mutant.

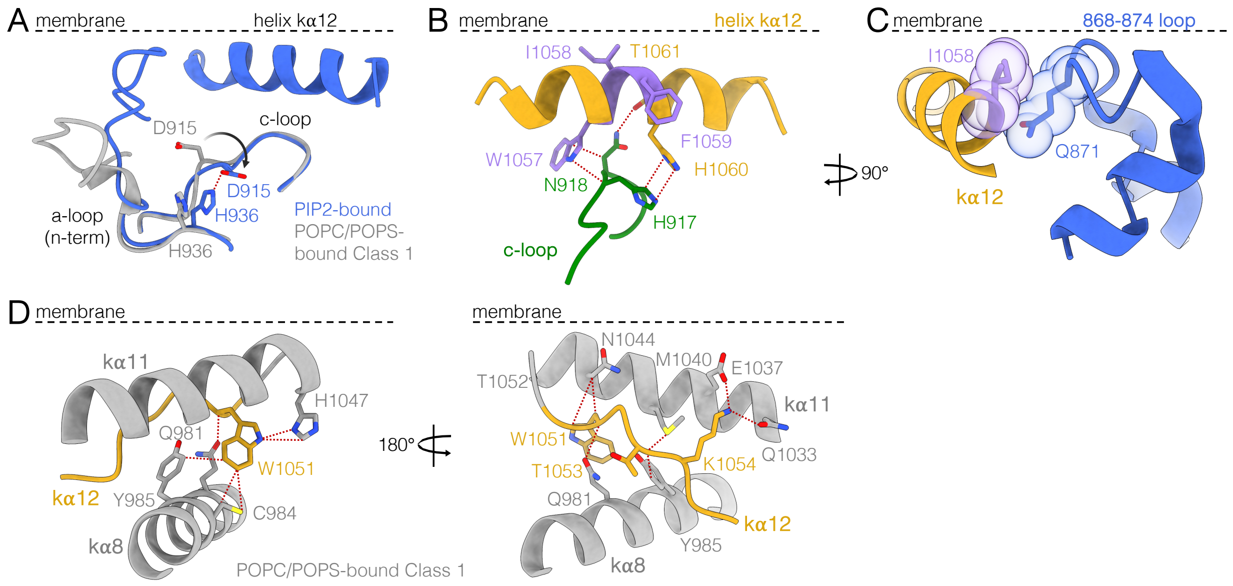


**Fig. S9. p110α active site with and without PIP2.** (**A**) POPC/POPS class 1 (gray) and PIP2-bound (blue) overlayed active sites, depicting the changing interaction between the catalytic loop (c-loop) and activation loop (a-loop) upon PIP2 binding. (**B**) Interactions between helix kα12 (yellow) and the c-loop (green) in the PIP2-bound state. WIF motif of helix kα12 is colored in purple. (**C**) Stabilization of membrane-interacting loop 868-874 by helix kα12 in the PIP2-bound complex. (**D**) Stabilization of helix kα12 in the inactive conformation by helices kα8 and kα11 in the POPC/POPS class 1 state.

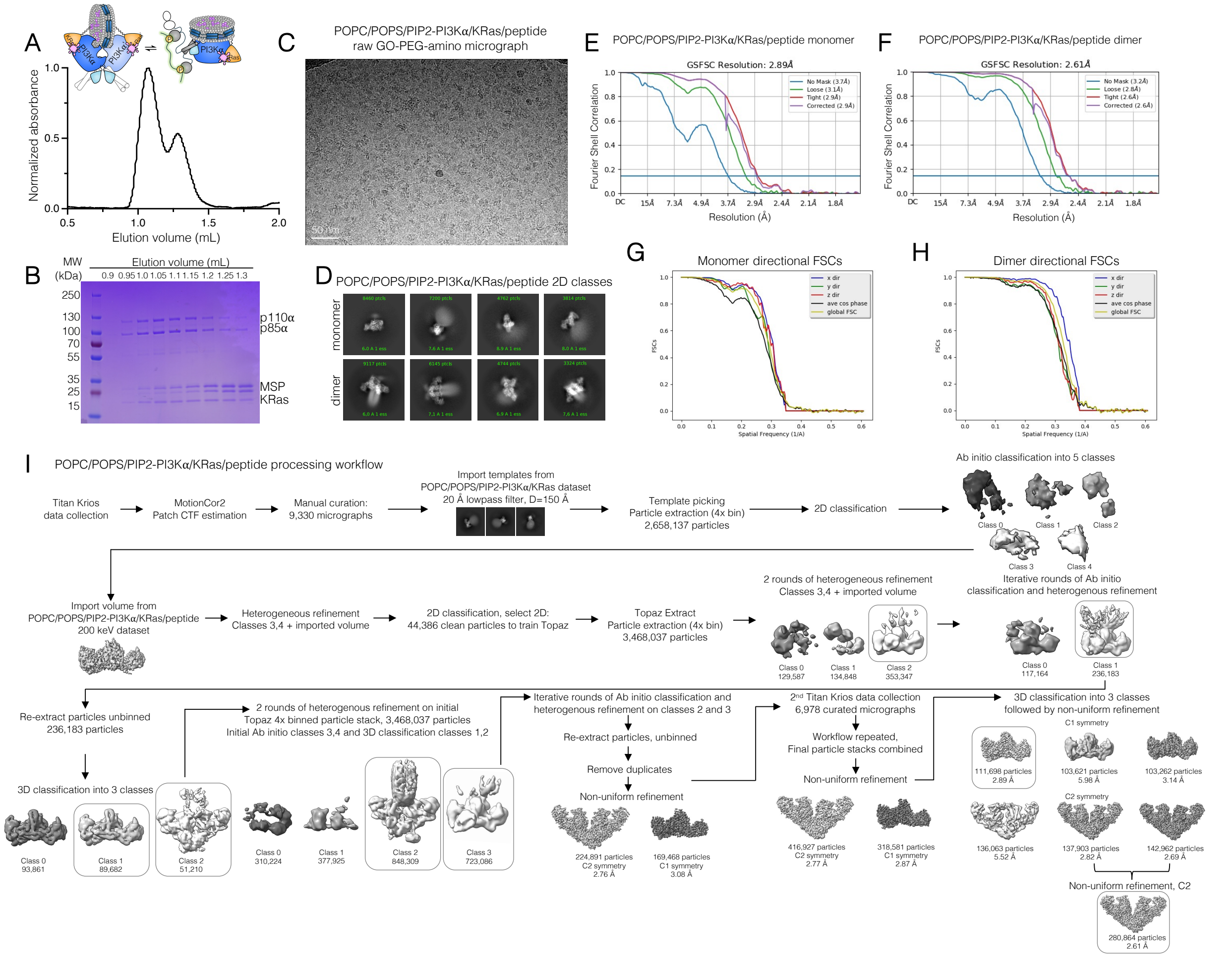


**Fig. S10. In-vitro reconstitution, map quality and processing workflow for the POPC/POPS/PIP2-PI3Kα/KRas/peptide complex dataset.** (**A**) Representative size exclusion chromatography profile of PI3Kα/KRas/peptide monomeric and dimeric complexes on POPC/POPS/PIP2/MSP1E3D1 nanodiscs resolved on a Superdex 200 Increase 3.2/300 column and (**B**) corresponding Coomassie-stained SDS-PAGE analysis. (**C**) Representative micrograph of the POPC/POPS-bound PI3Kα/KRas/peptide complex sample on Quantifoil R1.2/1.3 300 mesh Au holey-carbon PEG-amine functionalized graphene oxide grids from a dataset of 16,308 micrographs. The scale bar corresponds to 50 nm. (**D**) Representative cryo-EM 2D class averages of particles corresponding to the POPC/POPS/PIP2-PI3Kα/KRas/peptide monomeric and dimeric complexes. (**E-F**) Gold Standard Fourier Shell Correlation (GSFSC) of final maps used for model building for monomeric (E) and dimeric (F) complexes from CryoSPARC 4. (**G-H**) Directional FSCs of the monomeric (G) and dimeric (H) POPC/POPS/PIP2-PI3Kα/KRas/peptide complexes calculated by the 3DFSC server. (**I**) Processing workflow of POPC/POPS/PIP2-PI3Kα/KRas/peptide complex dataset.

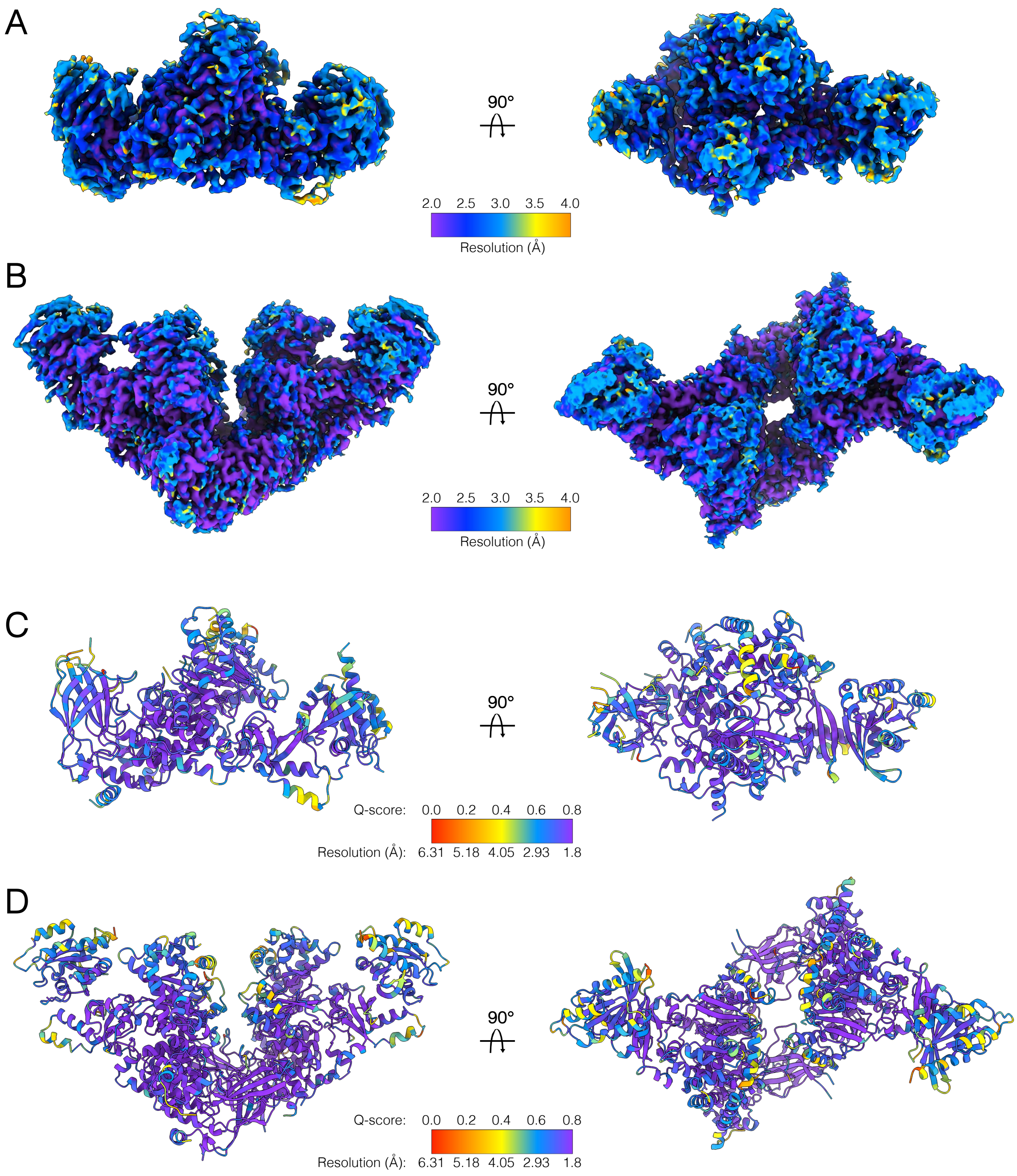


**Fig. S11. Resolution estimation and Q-score analysis of the POPC/POPS/PIP2-PI3Kα/KRas/peptide monomeric and dimeric complexes.** (**A-B**) Cryo-EM maps of the POPC/POPS/PIP2-bound PI3Kα/KRas monomeric (A) and dimeric (B) complexes colored according to local resolution determined by ResMap. (**C-D**) Corresponding models of POPC/POPS/PIP2-bound PI3Kα/KRas monomeric (C) and dimeric (D) complexes colored by estimated per residue Q-score. Color bars indicate corresponding resolution in Å (A-B) or estimated resolution in Å for reported Q-scores (C-D).

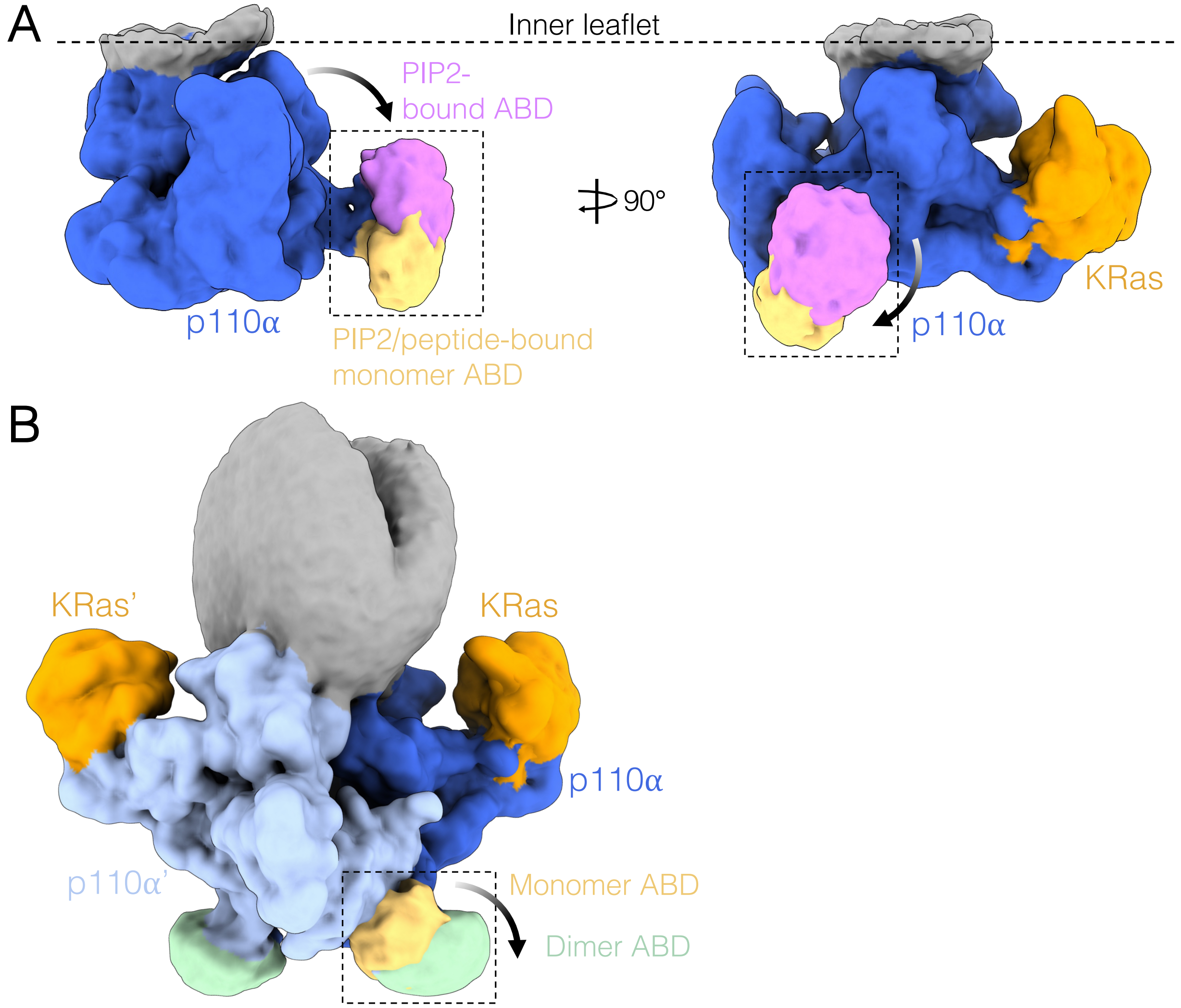


**Fig. S12. ABD displacement in PI3Kα activation.** (**A**) Overlayed PIP2-bound and PIP2/peptide-bound 8 Å low-pass filtered maps, depicting the gradual rotation of the ABD domain from the p110α catalytic core. (**B**) Overlayed PIP2/peptide-bound monomer and dimer maps, depicting the maximal rotation of the ABD domain from the p110α catalytic core in the dimer. p110α is shown in blue, KRas in orange, PIP2-bound ABD in pink, PIP2/peptide-bound monomer ABD in yellow and dimer ABD in green.

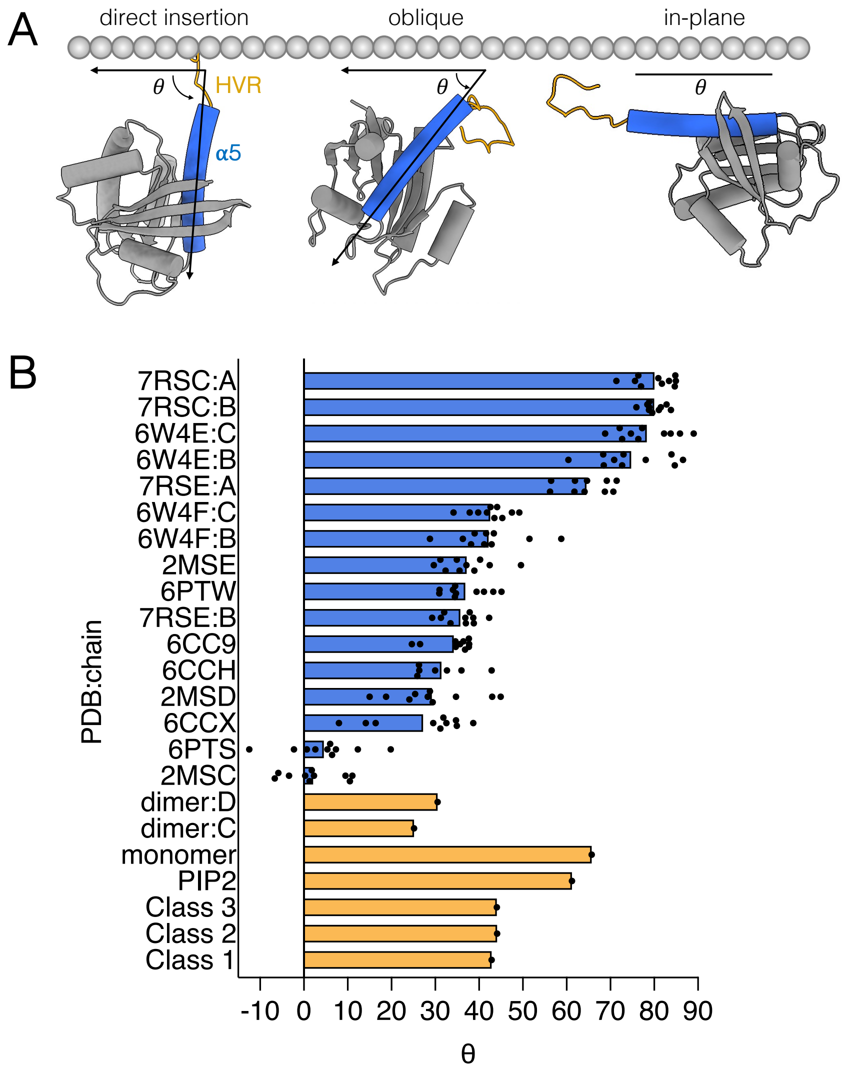


**Fig. S13. Dynamic membrane insertion of Ras.** (**A**) Examples of Ras membrane insertion angles from the Protein Data Bank (PDB). (**B**) Insertion angles of Ras as part of nanodisc-bound NMR structures from the PDB and KRas as part of our membrane-bound PI3K$\alpha$/KRas complex structures. Measurements represent the angle between helix $\alpha 5$ and the plane of the nanodisc. Each data point within a PDB corresponds to individual models within the NMR ensemble.

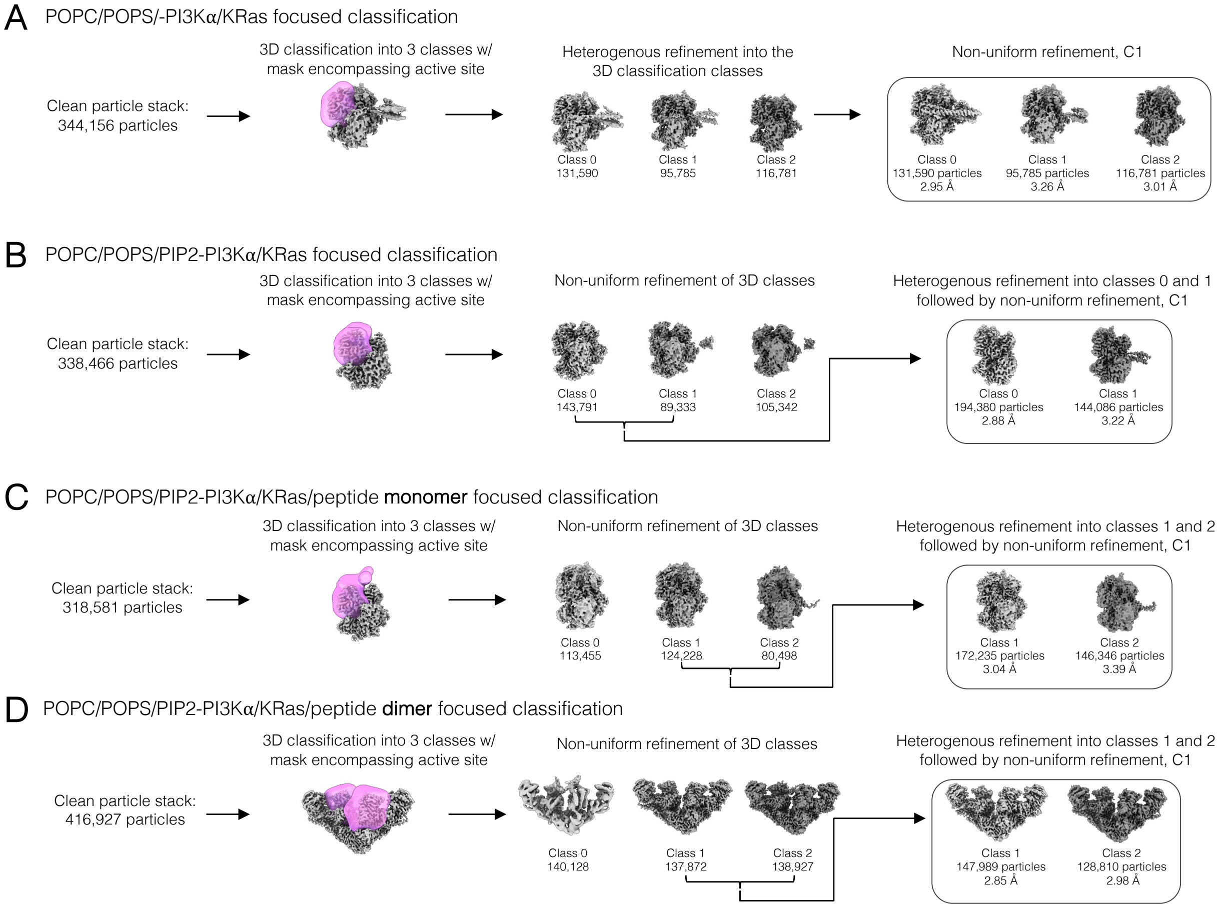


**Fig. S14. Processing workflow for focused classification of all datasets.** (**A-D**) Processing workflow of focused classification for (A) POPC/POPS-PI3Kα/KRas, (B) POPC/POPS/PIP2-PI3Kα/KRas, (C) POPC/POPS/PIP2-PI3Kα/KRas/peptide monomer and (D) dimer datasets. Focus masks are shown in pink. Final models are indicated with gray boxes.

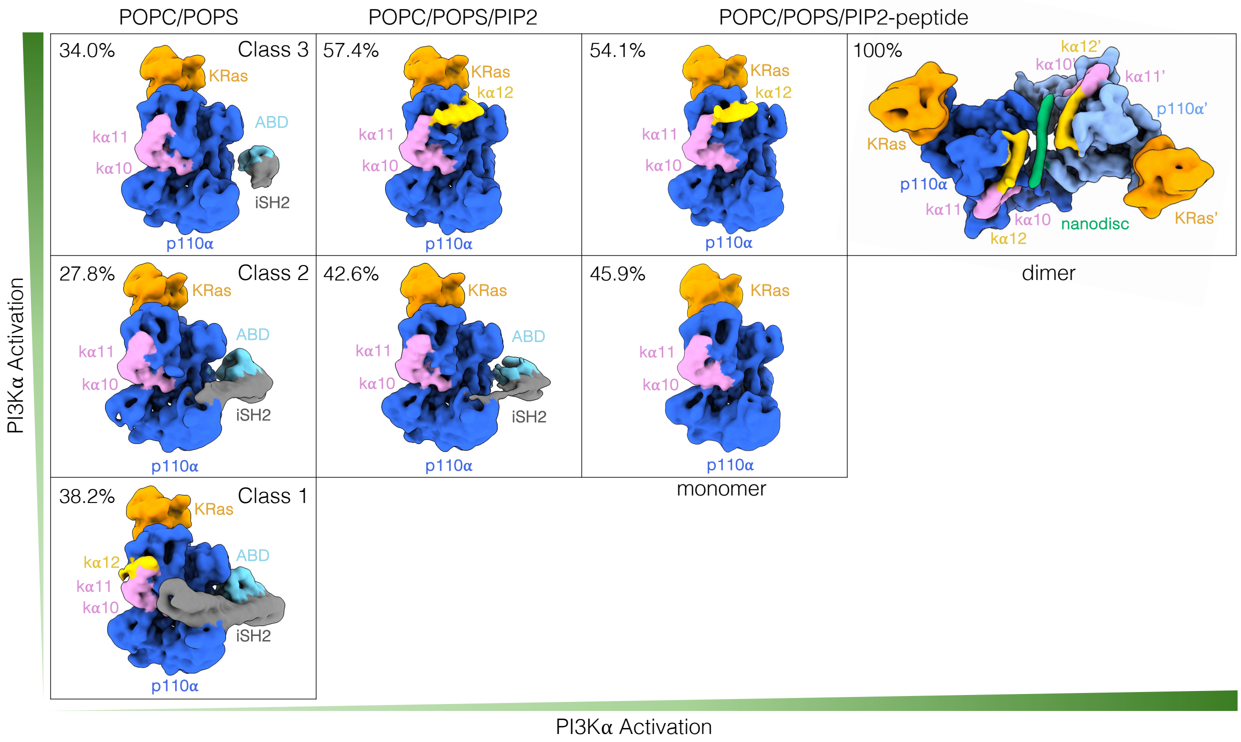


**Fig. S15. Allosteric communication between p85α and the p110α active site.** 8 Å low-pass filtered maps of final focused classification classes illustrating the release of inhibory iSH2 and ABD domains and the stabilization of helix kα12 in the active, inward conformation. The iSH2 domain of p85α is shown in gray, KRas in orange, p110α in blue, and the ABD, kα10 and kα11 helices, and helix kα12 in cyan, pink, and yellow respectively.

**Table S1. nSH2 and iSH2 states in deposited structures of autoinhibited PI3Kα**

| PDB ID | p85α construct | p110α construct | nSH2/iSH2 state |
|---|---|---|---|
| 8TSA | niSH2 | WT | ordered/ordered |
| 8TSD | niSH2 | WT | ordered/ordered |
| 5XGH | niSH2 | WT | ordered/ordered |
| 4YKN | niSH2 | WT | ordered/ordered |
| 8TS7 | niSH2 | WT | ordered/ordered |
| 5XGI | niSH2 | WT | ordered/ordered |
| 5XGJ | niSH2 | WT | ordered/ordered |
| 4L2Y | niSH2 | WT | ordered/ordered |
| 8TSB | niSH2 | WT | ordered/ordered |
| 4L1B | niSH2 | WT | ordered/ordered |
| 4L23 | niSH2 | WT | ordered/ordered |
| 5SWG | niSH2 | WT | ordered/ordered |
| 5SWR | niSH2 | WT | ordered/ordered |
| 5SXA | niSH2 | WT | ordered/ordered |
| 5SX8 | niSH2 | WT | ordered/ordered |
| 5SXJ | niSH2 | WT | ordered/ordered |
| 4OVU | niSH2 | WT | ordered/ordered |
| 5SXE | niSH2 | WT | ordered/ordered |
| 5SXF | niSH2 | WT | ordered/ordered |
| 5SXB | niSH2 | WT | ordered/ordered |
| 5SXI | niSH2 | WT | ordered/ordered |
| 2RD0 | niSH2 | WT | disordered/ordered |
| 5SXC | niSH2 | WT | ordered/ordered |
| 5SX9 | niSH2 | WT | ordered/ordered |
| 5SXD | niSH2 | WT | ordered/ordered |
| 5SWO | niSH2 | WT | ordered/ordered |
| 5SWP | niSH2 | WT | ordered/ordered |
| 5SW8 | niSH2 | WT | ordered/ordered |
| 5SXK | niSH2 | WT | ordered/ordered |
| 5SWT | niSH2 | WT | ordered/ordered |
| 6NCT | niSH2 | WT | ordered/ordered |
| 4OVV | niSH2 | WT | ordered/ordered |
| 7MYN | full-length | WT | ordered/ordered |
| 7MYO | full-length | WT | ordered/ordered |
| 8DCP | full-length | WT | ordered/ordered |
| 8TU6 | full-length | WT | ordered/ordered |
| 8ILV | full-length | WT | ordered/ordered |
| 8DCX | full-length | WT | ordered/ordered |
| 8DD4 | full-length | WT | ordered/ordered |
| 8TDU | full-length | WT | ordered/ordered |
| 8DD8 | full-length | WT | ordered/ordered |
| 8ILS | full-length | WT | ordered/ordered |
| 8ILR | full-length | WT | ordered/ordered |

**Table S2. Cryo-EM collection, refinement and resulting models statistcs**

| | POPC/POPS Class 1 (EMD-49454) (PDB 9NI6) | POPC/POPS Class 2 (EMD-49456) (PDB 9NI8) | POPC/POPS Class 3 (EMD-49455) (PDB 9NI7) | PIP2 (EMD-49451) (PDB 9NI3) | PIP2/peptide Monomer (EMD-49453) (PDB 9NI5) | PIP2/peptide Dimer (EMD-49452) (PDB 9NI4 ) |
|---|---|---|---|---|---|---|
| **Data collection and processing** | | | | | | |
| Magnification | 105,000x | 105,000x | 105,000x | 105,000x | 105,000x | 105,000x |
| Voltage (kV) | 300 | 300 | 300 | 300 | 300 | 300 |
| Total dose (e–/Å$^2$) | 47.7 | 47.7 | 47.7 | 47.7 | 47.7 | 47.7 |
| Dose rate (e-/physical pixel/sec) | 16 | 16 | 16 | 16 | 16 | 16 |
| Exposure per frame (sec) | 0.025 | 0.025 | 0.025 | 0.025 | 0.025 | 0.025 |
| Defocus range (µm) | -1.0 to -2.0 | -1.0 to -2.0 | -1.0 to -2.0 | -1.0 to -2.0 | -1.0 to -2.0 | -1.0 to -2.0 |
| Pixel size (Å) | 0.8189 (physical) | 0.8189 (physical) | 0.8189 (physical) | 0.8189 (physical) | 0.8189 (physical) | 0.8189 (physical) |
| Symmetry imposed | C1 | C1 | C1 | C1 | C1 | C2 |
| Initial particle images (no.) | 2,870,914 | 2,870,914 | 2,870,914 | 4,088,012 | 6,259,852 | 6,259,852 |
| Final particle images (no.) | 111,772 | 114,960 | 117,424 | 223,376 | 111,698 | 280,864 |
| Map resolution (Å) FSC threshold (0.143) | 3.01 | 3.23 | 3.05 | 2.82 | 2.89 | 2.61 |
| Map resolution range (Å) | 2.0-4.0 | 2.5-5.0 | 2.0-3.5 | 2.0-3.5 | 2.0-3.5 | 2.0-3.0 |
| **Refinement** | | | | | | |
| Initial model used (PDB code) | AF-P42336-F1-v4 AF-P27986-F1-v4 AF-P01116-F1-v4 | AF-P42336-F1-v4 AF-P27986-F1-v4 AF-P01116-F1-v4 | AF-P42336-F1-v4 AF-P01116-F1-v4 | AF-P42336-F1-v4 AF-P01116-F1-v4 | AF-P42336-F1-v4 AF-P01116-F1-v4 | AF-P42336-F1-v4 AF-P01116-F1-v4 |

| | POPC/POPS Class 1 (EMD-49454) (PDB 9NI6) | POPC/POPS Class 2 (EMD-49456) (PDB 9NI8) | POPC/POPS Class 3 (EMD-49455) (PDB 9NI7) | PIP2 (EMD-49451) (PDB 9NI3) | PIP2/peptide Monomer (EMD-49453) (PDB 9NI5) | PIP2/peptide Dimer (EMD-49452) (PDB 9NI4 ) |
|---|---|---|---|---|---|---|
| Model resolution (Å)<br>FSC threshold 0.5 (Masked) | 3.1 | 3.4 | 3.1 | 2.9 | 3.0 | 2.7 |
| Map sharpening *B* factor (Å$^2$) | -103.5 | -111.9 | -103.3 | -108.3 | -101.9 | -101.5 |
| Model composition<br>Non-hydrogen atoms<br>Protein residues | 10948<br>1321 | 9806<br>1189 | 8730<br>1063 | 8962<br>1089 | 8881<br>1081 | 17917<br>2180 |
| *B* factors (Å$^2$)<br>Protein | 49.81 | 107.76 | 34.98 | 27.52 | 34.18 | 26.65 |
| R.M.S. deviations<br>Bond lengths (Å)<br>Bond angles (°) | 0.014<br>1.927 | 0.013<br>1.867 | 0.015<br>2.014 | 0.014<br>2.022 | 0.015<br>2.021 | 0.014<br>1.968 |
| Validation<br>MolProbity score<br>Clashscore<br>Poor rotamers (%) | 0.79<br>0.96<br>0.82 | 0.77<br>0.87<br>0.55 | 0.55<br>0.12<br>0.31 | 0.66<br>0.45<br>0.81 | 0.59<br>0.23<br>0.71 | 0.59<br>0.22<br>0.25 |
| Ramachandran plot<br>Favored (%)<br>Allowed (%)<br>Disallowed (%) | 98.32<br>1.68<br>0.00 | 98.37<br>1.63<br>0.00 | 98.09<br>1.91<br>0.00 | 98.15<br>1.85<br>0.00 | 98.22<br>1.78<br>0.00 | 98.10<br>1.90<br>0.00 |

**Table S3. Refinement and resulting statistcs of low resolution models**

| **Refinement** | 10 Å LPS POPC/POPS Class 1 (EMD-49519) (PDB 9NLC) | 5 Å LPS PIP2 (EMD-49460) (PDB 9NIF) | 5 Å LPS PIP2/peptide Monomer (EMD-49459) (PDB 9NIE) | 5 Å LPS PIP2/peptide Dimer (EMD-49458) (PDB 9NID) |
|---|---|---|---|---|
| Initial model used (PDB code) | AF-P42336-F1-v4<br>AF-P27986-F1-v4<br>AF-P01116-F1-v4 | AF-P42336-F1-v4<br>AF-P01116-F1-v4 | AF-P42336-F1-v4<br>AF-P01116-F1-v4 | AF-P42336-F1-v4<br>AF-P01116-F1-v4 |
| Model resolution (Å)<br>FSC threshold 0.5 (Masked) | 6.2 | 3.0 | 3.1 | 2.9 |
| Map sharpening *B* factor (Å$^2$) | N/A | N/A | N/A | N/A |
| Model composition<br>Non-hydrogen atoms<br>Protein residues | 13058<br>1581 | 9064<br>1100 | 8973<br>1091 | 18105<br>2200 |
| *B* factors (Å$^2$)<br>Protein | N/A | N/A | N/A | N/A |
| R.M.S. deviations<br>Bond lengths (Å)<br>Bond angles (°) | 0.013<br>1.680 | 0.014<br>2.025 | 0.015<br>1.996 | 0.014<br>1.964 |
| Validation<br>MolProbity score<br>Clashscore<br>Poor rotamers (%) | 1.07<br>1.73<br>0.48 | 0.67<br>0.50<br>0.60 | 0.62<br>0.34<br>0.60 | 0.63<br>0.28<br>0.15 |
| Ramachandran plot<br>Favored (%)<br>Allowed (%)<br>Disallowed (%) | 97.26<br>2.74<br>0.00 | 98.35<br>1.65<br>0.00 | 98.05<br>1.95<br>0.00 | 97.89<br>2.11<br>0.00 |
